# Loss of MLL3/4 decouples enhancer H3K4 monomethylation, H3K27 acetylation, and gene activation during ESC differentiation

**DOI:** 10.1101/2022.10.24.513607

**Authors:** Ryan M. Boileau, Kevin X. Chen, Robert Blelloch

## Abstract

Enhancers are essential in defining cell fates through the control of cell type specific gene expression. Enhancer activation is a multi-step process involving chromatin remodelers and histone modifiers including the monomethylation of H3K4 (H3K4me1) by MLL3 (KMT2C) and MLL4 (KMT2D). MLL3/4 are thought to be critical for enhancer activation and cognate gene expression including through the recruitment of acetyltransferases for H3K27. Here we test this model by evaluating the impact of MLL3/4 loss on chromatin and transcription during early embryonic stem cell differentiation. We find that MLL3/4 activity is required at most if not all sites that gain or lose H3K4me1 but is largely dispensable at sites that remain stably methylated during this transition. This requirement extends to H3K27 acetylation (H3K27ac) at most transitional sites. However, many sites gain H3K27ac independent of MLL3/4 or H3K4me1 including enhancers regulating key factors in early differentiation. Furthermore, despite the failure to gain active histone marks at thousands of enhancers, transcriptional activation of nearby genes is largely unaffected, thus uncoupling the regulation of these chromatin events from transcriptional changes during this transition. These data challenge current models of enhancer activation and imply distinct mechanisms between stable and dynamically changing enhancers. Collectively, our study highlights gaps in knowledge about the steps and epistatic relationships of enzymes necessary for enhancer activation and cognate gene transcription.

## Background

Gene regulatory networks that drive cell fate are regulated spatiotemporally by cell type specific transcription factors (TFs). The critical functions of TFs in development are coupled to their target genes through TF binding of *cis*-regulatory elements such as enhancers. Target gene regulation is mediated by general chromatin regulators which are recruited to enhancers bound by TFs. As such, chromatin regulators are an essential part of enhancer function and their dysregulation has severe consequences in development and disease. For example, MLL3 and MLL4 are functionally redundant histone methyltransferases within the COMPASS complex which deposit the histone modification H3K4me1 primarily at enhancers(1, 2). Moreover, mutations in MLL3 and MLL4 cause developmental defects such as Kabuki Disease and are in the top ten of frequently mutated genes in cancer(3–5). Despite the importance of chromatin regulators to human health, we lack fundamental insight into how general chromatin regulators such as MLL3/4 coordinate enhancer function.

The interdependence of different histone modifications and their depositing enzymes at enhancers has been widely studied, although their mechanistic roles remain controversial. A prevailing model of enhancer activation is that DNA bound TFs recruit MLL3/4 which deposit H3K4me1(6–9). MLL3/4 in turn recruits the acetyltransferases P300/CBP which deposit H3K27ac. An activated enhancer then promotes expression of its cognate gene, most often one that is a nearby neighbor(10, 11). However, it is unclear how generalizable this model is. Indeed, recent studies have begun to challenge the importance of histone modifications and even chromatin regulators in gene regulation by enhancers(12–16).

Therefore, we decided to revisit the prevailing model including the epistatic relationship between MLL3/4 to H3K4me1, H3K27ac, and gene transcription in the context of ESCs transitioning from the naive to formative pluripotent states. The naive to formative transition is a rapid, homogenous, and well-characterized transition that faithfully recapitulates the early embryonic cell fate transition in epiblast cells of the mouse between E4.5 and E5.5 at the transcriptional and epigenomic level(Fig1A)(17–20). Importantly, this developmental window immediately precedes previously described cell fate specification and migrational phenotypes seen in gastrulation with the loss of MLL3/4 *in vivo*(6, 21). Therefore, the naive to formative transition provides an ideal model to study relationships in enhancer mechanics and transcriptional regulation.

Using state-of-the-art epigenomics techniques to profile genetic deletion of both MLL3 and MLL4 in the naive to formative transition we make several surprising discoveries. First, MLL3/4 is only required at a small subset of sites that maintain stable H3K4me1 levels during the transition but is essential for all sites that gain or lose H3K4me1. This demonstrates the existence of distinct mechanisms of H3K4me1 deposition at most stable vs dynamic sites. Second, loss of MLL3/4 reduces H3K27ac levels at most dynamic sites, but often does so independent of underlying H3K4me1 levels. Finally, despite dramatic loss of active histone modifications at many enhancers in the formative state, few loci can be linked to changes in nearby gene expression with these changes being relatively minor. Taken together, these results demonstrate that MLL3/4 protein and H3K4me1 deposition can be uncoupled from H3K27ac deposition and also from changes in gene expression at most enhancer sites during early ESC differentiation. Therefore, our data challenge the prevailing model of enhancer activation and suggests that the functional role of MLL3/4 protein in cell fate transitions is more context dependent than previously appreciated.

## Results

### MLL3/4 is dispensable for transcriptional activation of much of the formative program

To study the role of MLL3 and MLL4 in stem cell self-renewal and differentiation we acquired MLL3^-/-^; MLL4^fl/fl^ ESCs (hereafter called MLL3KO)(22). We integrated Cre-ERT2 into the *Rosa26* locus and induced Cre-recombination with tamoxifen to generate MLL3^-/-^; MLL4^-/-^ double knockout clones (DKO)(Fig.S1A). Western blotting confirmed loss of MLL3 protein alone in the parental line and both MLL3 and MLL4 in the DKO cells compared to a wildtype (WT) control (Fig.1B). Under naive pluripotency culture conditions (Serum+LIF+2i), the DKO cells formed colonies that were less compact and grew at a slower rate compared to their WT or MLL3KO counterparts (Fig.1C, Fig.S1B). Next, we removed LIF+2i from the media to induce the formative state after ∼63 hours in WT cells (23). As expected, WT ESC colonies flattened out during this transition. Similar phenotypic changes were seen in MLL3KO and DKO, although in both cases cells appeared more dispersed. Proliferation was reduced in MLL3KO and DKO cells relative to WT, with the greatest impact on transitioning DKO (Fig.S1B). All cell lines expressed OCT4 and NANOG protein in naive culture conditions and similarly downregulated NANOG in formative conditions as indicated by Western blot (Fig.S1C). These data show that while there are morphological and proliferative differences, the knockout cells self-renew under naive culture conditions, have a pluripotent-like cell identity, and show evidence of a cell state transition in formative conditions.

**Figure 1:**
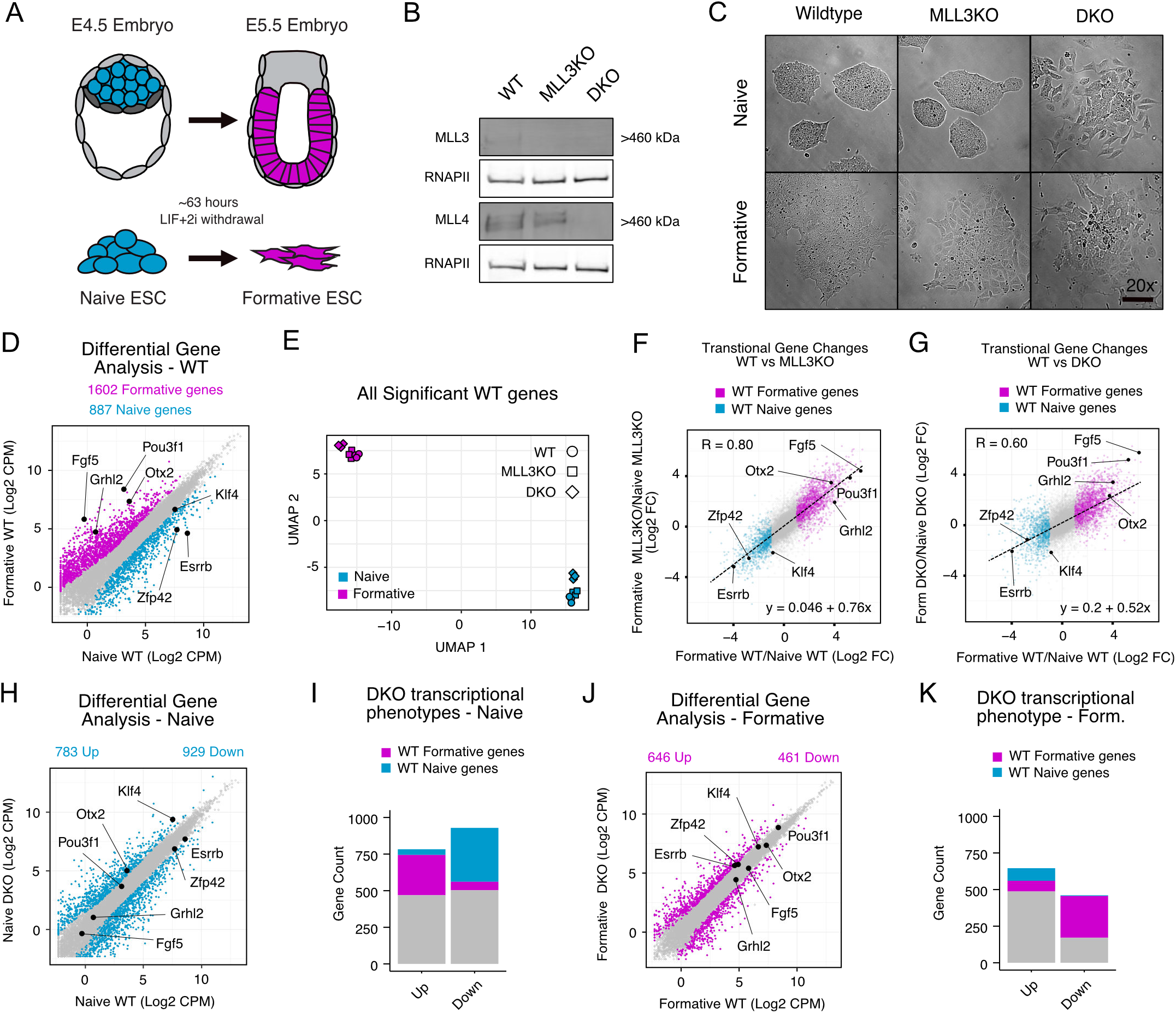
MLL3/4 is dispensable for transcriptional activation of much of the formative program. A) Schematic of naïve to formative transition *in vivo* and *in vitro*. B) Western validation of MLL3 and MLL4 knockout cell lines. C) Brightfield microscopy of cell lines in naive and formative conditions, 20x, scale bar = 100um. D) Differential gene expression analysis (DGE) on WT naive and formative RNA-seq samples (significant genes colored, P.adj < 0.05, Log2 Foldchange (Log2FC) >1). E) UMAP analysis on all RNA-seq samples subset by all naive and formative genes from WT DGE. F,G) Fold change (formative/naive) of all genes for either MLL3KO and DKO compared to WT. WT formative and naive genes from WT DGE analysis colored. R, Pearson’s coefficient. Dashed line and linear equation represent linear model of all genes. H) DGE analysis on WT naive and DKO naive samples (significant genes colored, P.adj < 0.05 and Log2FC > 1). I) Naive state DKO misexpressed genes categorized by whether they are also WT naive or formative genes. J,K) Same as H,I respectively but comparing within the formative state.

To determine the transcriptional effects of loss of MLL3/4 we performed RNA-seq on WT, MLL3KO, and DKO lines in naive and formative conditions. Differential gene expression analysis between WT naive and formative samples uncovered 887 downregulated genes (naive enriched genes) and 1602 upregulated genes (formative enriched genes) (Fig.1D). After subsetting for all naive and formative enriched genes we conducted UMAP analysis of both the WT and mutant cells under both conditions and found samples separated predominately by developmental state rather than by genotype (Fig. 1E). Correlation analysis of the gene expression changes seen in MLL3KO vs WT and DKO vs. WT showed Pearson correlations of 0.8 and 0.6 respectively showing that transitional changes in gene regulation still occur in the absence of MLL3/4 (Fig.1F,G). The dynamic range of gene expression changes was reduced in the MLL3/4 DKO as evidenced by a reduced slope in the linear model. This reduction was largely driven by diminished expression of naïve enriched genes in the naïve state (Fig.S1D). Upregulation of the formative program was largely unimpacted by MLL3/4 loss (Fig.S1D). qPCR analysis validated the successful upregulation of known formative markers (Fgf5, Otx2, Dnmt3b)(Fig.S1E). These results suggested a relatively normal overall transition in gene expression changes between naive and formative even in the absence of both MLL3 and MLL4.

In contrast, direct comparison of MLL3/4 DKO vs. WT identified hundreds of misregulated genes in both the naive and formative states. In the naive state, DKO cells had 783 upregulated genes and 929 downregulated genes (P.adj < 0.05 and Log2CPM > 1)(Fig.1H). We categorized these genes that were up or down by whether they are normally naive enriched, formative enriched, or unchanged between naive and formative (Fig.1I). This analysis showed a similar proportion of formative genes to be prematurely expressed (275 of 783, 35%) as naive enriched genes that were reduced in the naive state (348 of 928, 39%). In the formative state we found 646 genes were upregulated and 461 genes were downregulated in DKO cells compared to WT(Fig.1J). Surprisingly, fewer genes were down in the DKOs in the formative state than the naive state (461 vs. 929). Focusing on genes that normally change during the transition, 83 naive enriched genes were abnormally up and 283 formative enriched genes were abnormally down in the formative state (Fig.1K). These numbers were overall low relative to normal gene expression changes seen with naive to formative transition (887 down, 1602 up) showing that MLL3/4 only regulates a minor subset of genes among those that are normally gained with the transition from naive to formative. Given that the WT and DKO ESCs represent slightly different strain backgrounds (see methods), we also compared MLL3/4DKO to its MLL3KO parental line. This comparison showed even fewer impacted genes (Fig.S1F,G).

### MLL3/4 is required for all dynamic H3K4me1 deposition during pluripotent transition

Though expression of a small number of genes were impacted it was unclear if this was due to direct or indirect effects of MLL3/4 loss. To gain a better understanding of the direct effects of MLL3/4 loss on transcription changes during the naive to formative transition we evaluated the monomethylation of H3K4, the primary known role of MLL3/4 and a surrogate marker of their binding. We performed CUT&RUN(24) to profile H3K4me1 in naive and formative samples for WT, MLL3KO, and DKO cells. Differential peak analysis using DiffBind(25) on WT naive and formative samples with peaks called by SEACR(26) uncovered approximately ∼13000 peaks each that significantly gained (formative H3K4me1 peaks) or lost H3K4me1 (naive H3K4me1 peaks) during the normal pluripotent transition (FDR < 0.01, Log2 FC > 1). An additional ∼94000 peaks did not significantly change between cell states (shared H3K4me1 peaks, FDR > 0.1, Log2 FC < 0.7)(Fig.2A,B). Naïve and formative enriched peaks were almost completely lost in the DKO cells, but only slightly reduced in MLL3KO cells (Fig.2C,D). Reanalysis by Diffbind uncovered 0 total peaks that showed a significant change between the two states in the DKO background while we found 29088 significantly changing peaks in MLL3KO compared to 27458 total peaks in WT (Fig.2E, Fig.S2A). In contrast, shared H3K4me1 peaks (shown in gray) were not obviously affected. To test whether the loss of dynamic H3K4me1 was a direct effect of MLL3/4 enzymatic activity, we performed and evaluated RNA-seq and H3K4me1 CUT&RUN data using ESCs with point mutations in the SET domains of both MLL3 and 4 that render them catalytically dead (dCD cells)(12). Consistent with previous reports(12, 27), we found a comparable reduction in global H3K4me1 in westerns of acid-extracted histones between DKO and dCDs when compared to WT while seeing few transcriptional differences between dCD and WT cells in the naive and formative states (Fig.S2B-E). Moreover, H3K4me1 CUT&RUN on dCD cells largely recapitulated the H3K4me1 defects observed in DKOs (Fig.2F). Diffbind uncovered only 272 total differential peaks between naive and formative in the catalytic mutant background (Fig.2G).

**Figure 2:**
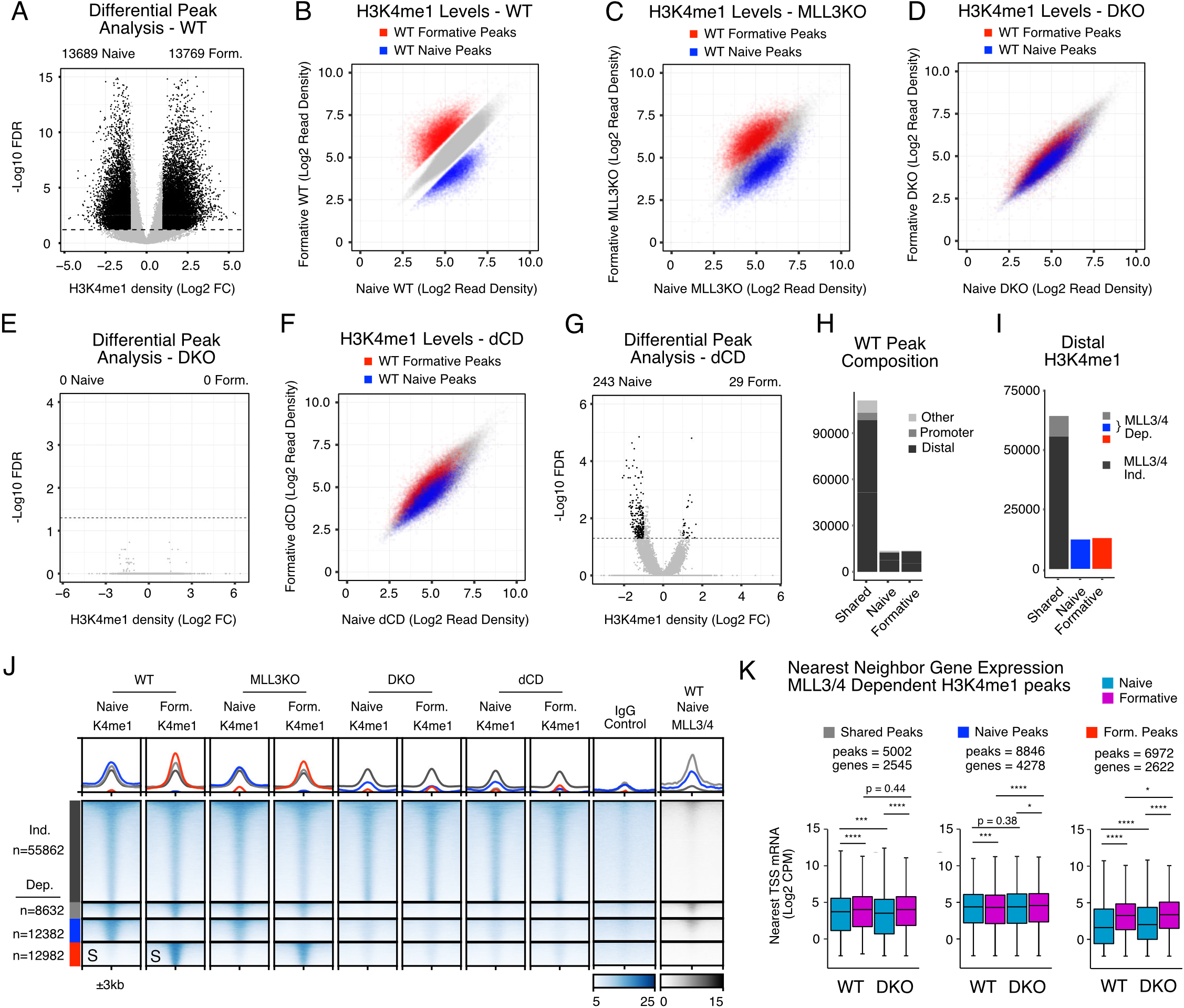
MLL3/4 is required for all dynamic H3K4me1 deposition during pluripotent transition. A) Differential signal enrichment between naive and formative WT H3K4me1 peaks identifies significant peaks (Black, FDR <0.05, Log2 Foldchange (Log2FC) > 1). B) Scatterplot of H3K4me1 peak intensities between naive and formative WT samples. (gray, shared peaks, FDR> 0.1 & Log2FC < 0.7, red or blue, FDR < 0.05 & Log2FC > 1). C,D) H3K4me1 signal in naive and formative MLL3KO or DKO samples at peaks called from WT. Colors correspond to peak categories derived from WT. E) Differential signal enrichment of naive and formative DKO samples. F) Same as C,D but with naive and formative dCD cells. G) Differential signal enrichment of naive and formative dCD samples. H) Feature annotation of WT peaks stratified by peak category including intergenic/intronic (Distal), Promoter, and all else (Others). I) Distal H3K4me1 peak categories further stratified by MLL3/4 independent (DKO/WT Log2FC > -0.7) or dependent (DKO/WT Log2FC < -1). J) Heatmap of MLL3/4 independent and dependent H3K4me1 peak categories, rows sorted on “S” columns. All heatmap values and range are in CPM. For metagene analysis the range in CPM is the same as shown in heatmap for each factor. K) nearest neighbor TSS analysis of expression levels in Log2CPM for each RNA-seq dataset near H3K4me1 peak categories. Multi-comparison paired Wilcoxon Rank-Sum Test, Benjamini-Hochberg corrected. *p<0.05 **p<0.01 ***p<0.001 ****p<0.0001

The majority of H3K4me1 sites, whether shared, naive-enriched, or formative-enriched were located at distal elements (intergenic or intronic) and not promoters (Fig.2H). To quantify dependency of MLL3/4 H3K4me1 at the various sites, we set cutoffs for dependent (Log2 FC naive DKO/WT < -1 or Log2 FC formative DKO/WT < -1) and independent H3K4me1 (Log2 FC naive DKO/WT > 0.7 & Log2 FC form DKO/WT < 0.7). Of the distal H3K4me1 sites, only 15% of shared sites (8632 of 64494) showed dependency on MLL3/4 for H3K4me1, while all naive and formative specific sites were dependent. Heatmap visualization validated these categories and similar behavior at dynamic sites during the transition was observed with analysis of published naive and formative H3K4me1 ChIP-seq datasets (Fig.2J and Fig.S2F)(23, 28). Furthermore, evaluation of published naive MLL3/4 ChIP-seq data showed enrichment of the proteins at H3K4me1 sites whether naïve-enriched or shared (12). Together, these data show that MLL3/4 are essential for a subset of H3K4me1 peaks including all dynamic sites and a small fraction of shared sites.

Next, we asked about the impact of MLL3/4 loss on the transcription of genes closest to the sites representing the different MLL3/4 dependent categories (shared, naive, and formative H3K4me1 sites) (Fig. 2K). Genes near MLL3/4 dependent shared H3K4me1 sites showed a clear upregulation in expression in the transition from naive to formative in WT cells, suggesting these enhancers may be primed in the naive state and became activated in the formative state. The loss of MLL3/4 had no impact on the upregulation of these genes with the transition to the formative state, although there was a slight reduction in expression in the naïve state. This later finding is consistent with previous findings showing downregulation of genes near MLL3/4 peaks in steady state naïve DKO ESCs(12), something we also saw with our expression data (Fig.S2G). In contrast, genes nearby MLL3/4 dependent naive-specific H3K4me1 sites were not impacted in the naïve DKO ESCs. Most surprising, while genes nearby MLL3/4 dependent formative-specific H3K4me1 showed a strong gain in expression during the naive to formative transition, this gain was not diminished by the loss of MLL3/4. Together, these results show that while MLL3/4 is required for all dynamic H3K4me1 and a subset of shared H3K4me1 between the naive and formative state, this requirement has minimal impact on gain of nearby gene transcription with the transition to the formative state.

### MLL3/4 dependent and independent distal H3K27ac deposition

MLL3/4 is thought to be important for the recruitment of histone acetyltransferase P300/CBP and thus histone acetylation including at H3K27 at enhancers. To test this general model at the genomic level, we conducted CUT&TAG(29) for H3K27ac in WT and DKO cells in the naive and formative states. We performed Diffbind at all peaks called by SEACR, intersected these with ATAC-seq peaks from naive and formative WT cells, and subset for distal sites to further enrich for active enhancers. A total of 12802 H3K27ac peaks were significantly down (naive H3K27ac peaks) and 8924 peaks were significantly up (formative H3K27ac peaks) during the transition (Fig.3A,B). Another 12205 peaks did not show significant change in H3K27ac between the two cell states (shared H3K27ac peaks). Loss of MLL3/4 resulted in a dramatic decrease in the dynamic changes in H3K27ac normally seen in the WT cells (Fig. 3C). However, unlike with H3K4me1, many sites still showed some change including sites that appeared to be unique to each pluripotent state. Diffbind using DKO samples uncovered 4295 naive-enriched, 1935 formative-enriched, and 23454 shared peaks, a reduction compared to WT, but still suggestive of cell state dependent dynamic acetylation (Fig. 3D). In both WT or DKO cells, the majority of H3K27ac was located at distal sites, including both shared and naive/formative-enriched peaks. Most of the cell-state specific H3K27ac sites were dependent on MLL3/4 (Log2 FC DKO/WT < -1 or Log2 FC DKO/WT > -0.7 respectively) with MLL3/4 dependent sites accounting for 22% of shared (2661 of 12205), 57% at naive-enriched (5717 of 9963), and 62% at formative-enriched sites (4495 of 7205) (Fig.3F). Heatmap visualization confirmed these findings (Fig. 3G,H, Fig. S3A). Interestingly though, metagene analysis identified a low-level persistent signal and premature H3K27ac signal at MLL3/4 independent naive and formative distal sites respectively. Analysis of published ChIP-seq for H3K27ac and the acetyltransferase P300 in naive and formative cells, showed commensurate binding at the sites with correlated increases and decreases in signal with H3K27ac signal in WT cells (Fig.S3B)(23, 28). These data uncover both MLL3/4 dependent and independent mechanisms of P300 acetylation of H3K27 at cell-type specific enhancer sites.

**Figure 3:**
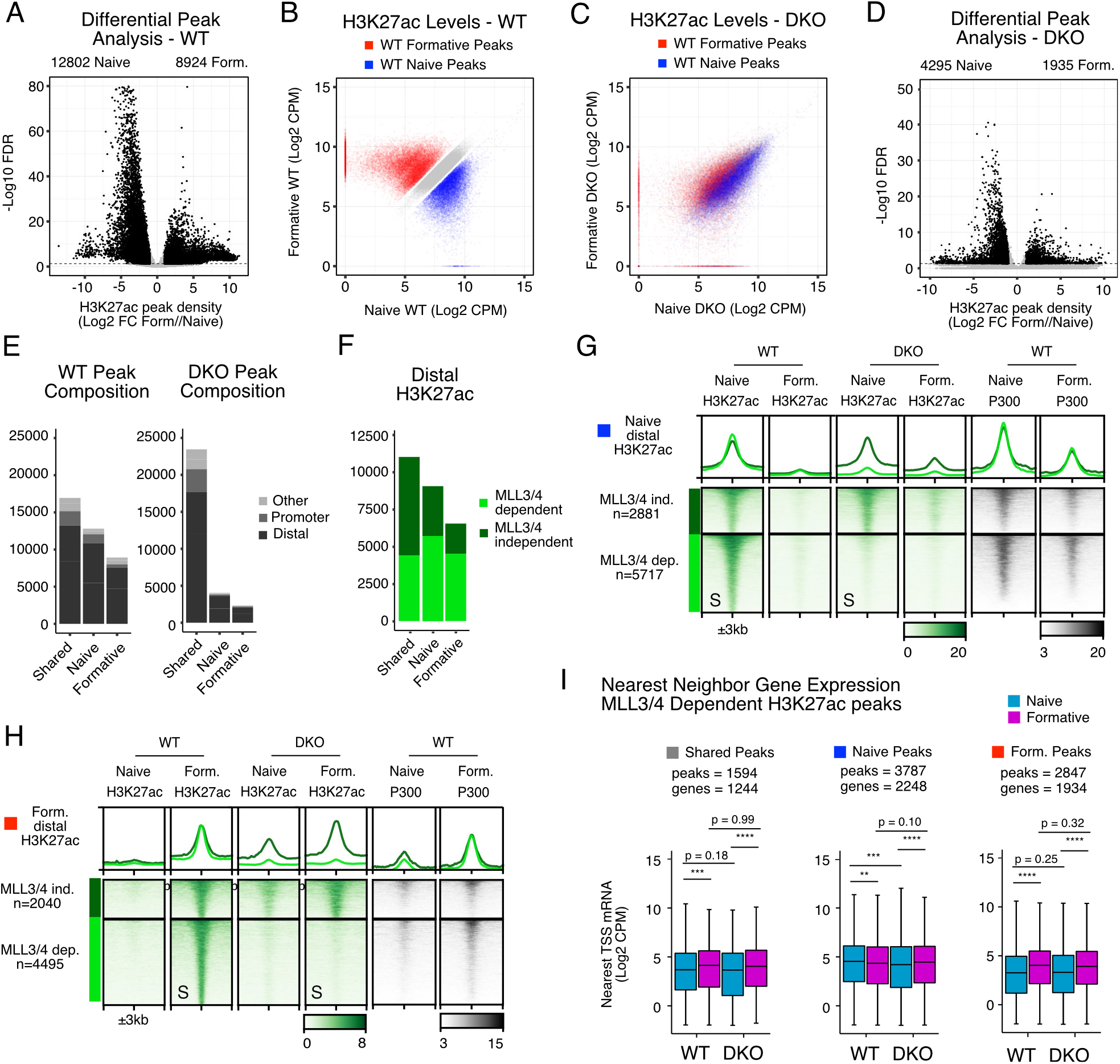
MLL3/4 dependent and independent distal H3K27ac deposition. A) Differential signal enrichment between naive and formative WT H3K27ac peaks identifies significant peaks (Black, FDR <0.05, Log2 Foldchange (Log2FC) > 1). B) Scatterplot of H3K27ac peak intensities between naive and formative WT samples. (gray, shared peaks, FDR > 0.1 & Log2FC < 0.7, red or blue, FDR < 0.05 & Log2FC > 1). C) H3K27ac signal in naive and formative DKO samples at peaks called from WT. Colors correspond to peak categories derived from WT. D) Feature annotation of WT or DKO peaks stratified by peak category including intergenic/intronic (Distal), Promoter, and all else (Others). E) Distal H3K27ac peak categories further stratified by MLL3/4 independent (DKO/WT Log2FC > -0.7) or dependent (DKO/WT Log2FC < -1). G,H) Heatmap of MLL3/4 independent and dependent H3K27ac peak categories for naive or formative specific peaks, rows sorted on “S” columns. All heatmap values and range are in CPM. For metagene analysis the range in CPM is the same as shown in heatmap for each factor. I) nearest neighbor TSS analysis of expression levels Log2CPM near H3K27ac peak categories. Multi-comparison paired Wilcoxon Rank-Sum Test, Benjamini-Hochberg corrected. *p<0.05 **p<0.01 ***p<0.001 ****p<0.0001

Given that H3K27ac is thought to be a marker of active enhancers, we next asked whether sites with MLL3/4 dependent H3K27ac had impacted transcription at neighboring genes upon loss of MLL3/4. We identified the nearest neighbor transcription start site (TSS) to all distal sites where H3K27ac was dependent on MLL3/4 and separated these sites by whether they were normally shared, naive-enriched, or formative-enriched. We then evaluated expression of genes associated with the TSSs (Fig. 3I). Shared peaks were associated with genes that normally increased in expression during the transition; this increase was not impacted by MLL3/4 loss. Naive-enriched peaks were associated with genes that show a slight reduction during the transition and that reduction occurred prematurely in the MLL3/4 DKO cells. Strikingly, there was a strong upregulation in gene expression nearby sites that normally show MLL3/4 dependent increases in H3K27ac, but this upregulation was not impacted by MLL3/4 loss. These data show that the upregulation of gene expression associated with a gain of H3K27ac at nearby enhancer sites is largely independent of that acetylation, while maintenance of gene expression associated with naive enriched H3K27ac sites is at least in part dependent on maintenance of that acetylation.

### Enhancer Activation Can Occur Independently of MLL3/4

The data above suggested similar trends in terms of the effects of MLL3/4 on H3K4me1, H3K27ac and nearby gene expression, consistent with a partial dependency on MLL3/4 recruitment for H3K27ac deposition. To investigate this dependency, we focused on all distal sites that show changes in H3K27ac in the context of their H3K4me1 status. We first separated all chromatin accessible H3K4me1 sites (ATAC+) into those that lose, gain, or have shared methylation during the naive to formative transition (Fig.S4A). Reanalyzing nearest neighbor transcription with these sites did not appreciably change our results obtained with all H3K4me1 sites (Fig.S4B vs. Fig.2K). Next, we subdivided the H3K4me1 categories into ones that lose, gain, maintain, or never have coincident H3K27ac (Fig.4A). In general, most H3K4me1 sites never showed H3K27ac. However, among the 5960 sites that lose H3K4me1 during the transition, 1370 also lose H3K27ac, and 913 maintain acetylation. Among 5583 sites that gain H3K4me1, 1172 sites also gain H3K27ac. We identified few sites (<150) for the remaining categories of H3K27ac at sites that either gain or lose H3K4me1. Among 37589 sites maintaining H3K4me1, 2264 sites gain acetylation, 3498 sites lose acetylation, 4353 sites show maintained acetylation. This suggests, although sites with dynamic H3K4me1 tended to have dynamic H3K27ac in the same direction (gain or loss), most H3K27ac dynamics during the naive to formative transition typically occur at sites with preexisting H3K4me1.

**Figure 4:**
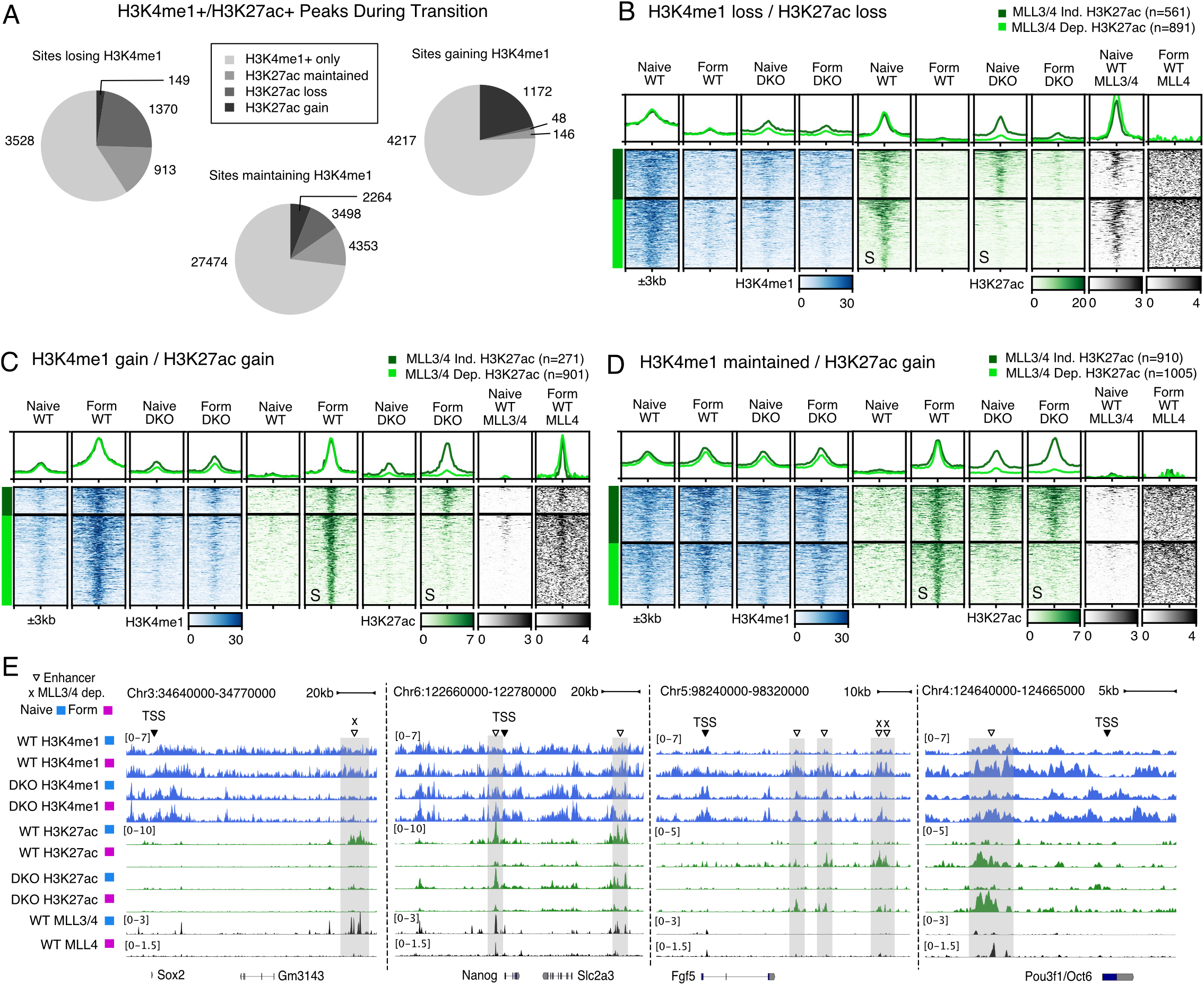
Enhancer Activation Can Occur Independently of MLL3/4. A) Pie charts showing fraction of coincident changes in H3K27ac within the different categories of changing H3K4me1 following MLL3/4. B) Heatmap of sites that normally lose H3K4me1 and H3K27ac during naive to formative transition clustered by either MLL3/4 dependent or independent H3K27ac. Rows sorted on “S” columns. Y-axis scale of metagene profiles above is same as signal ranges of heatmap, values in CPM. C) Same as B but using sites that normally gain H3K4me1 and H3K27ac during naive to formative transition clustered by either MLL3/4 dependent or independent H3K27ac. D) Same as B but using sites that have unchanging H3K4me1 but gain H3K27ac during naive to formative transition clustered by either MLL3/4 dependent or independent H3K27ac. All heatmap values and range are in CPM. For metagene analysis the range in CPM is the same as shown in heatmap for each factor. E) Naive and formative examples of functionally validated enhancers that are either MLL3/4 dependent or independent. Naive MLL3/4 dependent Sox2 enhancer cluster, Naive MLL3/4 independent Nanog enhancers, Formative independent and dependent enhancers of Fgf5, formative independent enhancer of Pou3f1/Oct6.

To understand how these subgroups were impacted by the loss of MLL3/4, we visualized the data as heatmaps and filtered for MLL3/4 independent or dependent H3K27ac (Log2FC DKO/WT > -0.7 or Log2FC DKO/WT < -1 respectively). Several important findings were uncovered by these efforts. Among the sites that are enriched for both H3K4me1 and H3K27ac (H3K4me1+/H3K27ac+) in the naive state, the loss of MLL3/4 resulted in a consistent loss of monomethylation as seen across all naive enriched H3K4me1 sites (Fig. 4B). However, only 61% of these sites lost H3K27ac (891 of 1452 sites). Among sites that gain H3K4me1 and H3K27ac during the transition from naive to formative, again all H3K4me1 is dependent on MLL3/4, while 77% also lost H3K27ac (901 out of 1172 sites)(Fig. 4C). To confirm that the naive and formative H3K4me1+/H3K27ac+ sites were direct MLL3/4 targets, we analyzed published naive ChIP-seq of MLL3/4 and generated CUT&RUN for MLL4 in the formative state (Fig. 4B,C). Naive MLL3/4 ChIP-seq signal was enriched at naive sites and MLL4 CUT&RUN was enriched specifically at formative sites. Together these analyses show H3K27ac maintenance and de novo deposition can occur at enhancer sites independent of MLL3/4 including at sites requiring MLL3/4 for H3K4me1 deposition.

Next, we analyzed the sites that normally retain monomethylation during the transition (H3K4me1 shared sites), which included a mix of sites that did or did not require MLL3/4 for H3K4me1 (Fig. 2J). Among the MLL3/4 independent H3K4me1 sites, 2039 sites normally gained H3K27ac, 2357 sites lost H3K27ac, and 3611 sites had H3K27ac that did not change. An additional 18135 sites never showed H3K27ac. For sites that gained H3K27ac, 48% required MLL3/4 to do so even though the same enzymes were not required for the monomethylation at those sites (Fig. 4D). The remaining 52% showed strong H3K27ac in formative cells independent of MLL3/4 and most showed premature H3K27ac deposition in naive DKO cells. MLL3/4 signal was low at all these sites consistent with the unaffected H3K4me1 which suggested the loss of acetylation at these sites was due to indirect effects. Similar to the formative H3K27ac sites, naive H3K27ac at shared H3K4me1 sites also demonstrated mixed dependency on MLL3/4 with only 844 of the sites losing H3K27ac (Fig.S4B). Among the remaining 1320 sites, which retained high levels of naive H3K27ac, the reduction in acetylation upon differentiation was incomplete in DKOs. In contrast to the formative sites, MLL3/4 signal was strong at the naive dependent sites. In MLL3/4 dependent shared H3K4me1 sites (2113 sites) few sites gained or had shared H3K27ac (225 and 742, respectively). Despite losing H3K4me1 these sites showed mixed dependency of H3K27ac on MLL3/4. Notably, sites with naive enriched H3K27ac comprised the largest group (1141 sites) and most of these sites (973 of 1141 sites, 85%) were dependent on MLL3/4 for H3K27ac. Finally, H3K4me1 independent shared sites showed no appreciable MLL3/4 binding despite a loss of H3K27ac in DKO cells at 992 of 3297 sites, implying indirect effects (Fig.S4D). Taken together, these data show that H3K27ac deposition at cell-type specific sites is not necessarily coupled with binding or activity of MLL3/4. Indeed, we identified almost as many exceptions as examples for the canonical model coupling MLL3/4 and H3K27ac deposition at enhancers.

Next, we asked how the loss of MLL3/4 impacted H3K4me1 and H3K27ac at well-known functionally validated enhancers regulating genes important for pluripotent cell identity including Sox2, Nanog, Fgf5, and Oct6(28,30–34). Sox2 and Nanog expression normally decrease, while Fgf5 and Oct6 expression increase during the naive to formative transition. In the MLL3/4 DKOs, Sox2 expression failed to decrease during the transition, while the other genes showed normal patterns of expression (Fig.1G, Fig.S4E). However, the impact of MLL3/4 loss on H3K4me1 and H3K27ac at associated enhancers for each of these genes varied (Fig. 4E). The Sox2 enhancer cluster showed stable H3K4me1 and naive specific H3K27ac, both of which were dependent on MLL3/4. Nanog has at least two known enhancers, all normally showing stable H3K4me1 and a loss of H3K27ac during the transition; none of which were impacted by MLL3/4 loss. Fgf5 has 4 known enhancers, all normally showing increases in H3K4me1 and H3K27ac during the transition. In the DKO cells, all four enhancers showed reduced H3K4me1, with two showing normal upregulation of H3K27ac and two showing reduced H3K27ac in the formative state. Oct6 has one known enhancer, which normally shows a gain in H3K4me1 and H3K27ac during the transition; H3K4me1 was reduced, but H3K27ac was unchanged in DKO cells. These examples of well-known enhancers demonstrate the variable nature of dependency on MLL3/4 for H3K4me1 and H3K27ac, including a surprising dispensability for de novo activation of enhancers for Fgf5 and Oct6 expression.

We hypothesized that dependency of H3K4me1/H3K27ac on MLL3/4 might vary based on the TFs bound at each enhancer. Therefore, we performed differential motif analysis using Gimmemotifs(35) to determine TF binding motifs enriched at enhancer sites dependent on MLL3/4 for H3K4me1 and/or H3K27ac (Fig.S5A). This analysis identified strong enrichment for the binding motifs for TCF7L2, GRHL, and ZIC1 at sites requiring MLL3/4 for the gain of both H3K4me1 and H3K27ac during the naive to formative transition. Expression of each of these TFs is maintained or increased during the transition, and GRHL2 is essential for the gain of H3K4me1 and H3K27ac at hundreds of formative specific enhancers(23). At MLL3/4 dependent naive peaks, we identified a strong enrichment for multiple TF binding motifs including RXRA, ZBTB20, ZNF257, MYC, ESR2, and NR6A1, all of which are either expressed or have closely related TFs that are expressed in naive cells (Fig.S5B). Interestingly, the motifs for the critical TF regulators of the naive and/or formative states such as Oct4, Sox2, Nanog, KLFs, Oct6, and Otx2, were generally not uncovered. These analyses suggest that TFs differ in their dependency on MLL3/4 to maintain or establish H3K27ac.

### Distal H3K4me1 and H3K27ac are not functionally coupled with formative transcriptional activation

We next asked whether focusing on sites of H3K27ac change in the context of dynamic H3K4me1 peaks might enrich for sites near genes whose expression was impacted by MLL3/4. Similar to analysis of H3K27ac alone, genes nearby naive H3K4me1+/H3K27ac+ sites with MLL3/4 dependent H3K27ac showed reduced expression in naive cells (Fig.4B,Fig.S6A) while genes nearby MLL3/4 independent H3K27ac were unchanged, supporting a role for acetylation in maintaining expression at these sites. In contrast, in the formative state there was little impact on transcription of genes nearby either MLL3/4 independent or dependent H3K27ac (Fig.4C,Fig.S6B). This suggested gain of both H3K4me1 and H3K27ac at formative sites was not associated with formative transcriptional activation.

To expand on this surprising finding, we used ChromHMM as an unbiased approach to look at all potential combinations of H3K4me1, H3K27ac in WT and DKO cells in the naive and formative states. We modeled 16 chromatin states to segment all possible combinations of these modifications, and subset for distal peaks overlapping WT ATAC peaks separately in naive and formative cells (Fig.5A,C). The sites were then mapped to nearby TSS, and the impact of MLL3/4 on expression of these genes was measured (Fig.5B,D). In the naive state, genes near sites that lost H3K27ac showed highly significant downregulation whether or not they also lost H3K4me1 (States 1,15). These sites also showed strong MLL3/4 signal supporting a direct role for the enzymes in coordinating both acetylation and transcription. In the formative state, only genes near distal sites that lost both H3K4me1 and H3K27ac showed a significant reduction in gene expression (State 2) (Fig. 5D). This state represented a relatively small number of enhancers (1040 sites) linked to only 154 genes. Therefore, this unbiased approach at defining chromatin states again showed that even though MLL3/4 loss dramatically alters the landscape of enhancer activity states, the impact on gene expression changes that occur during the naive to formative transition is relatively minor.

**Figure 5:**
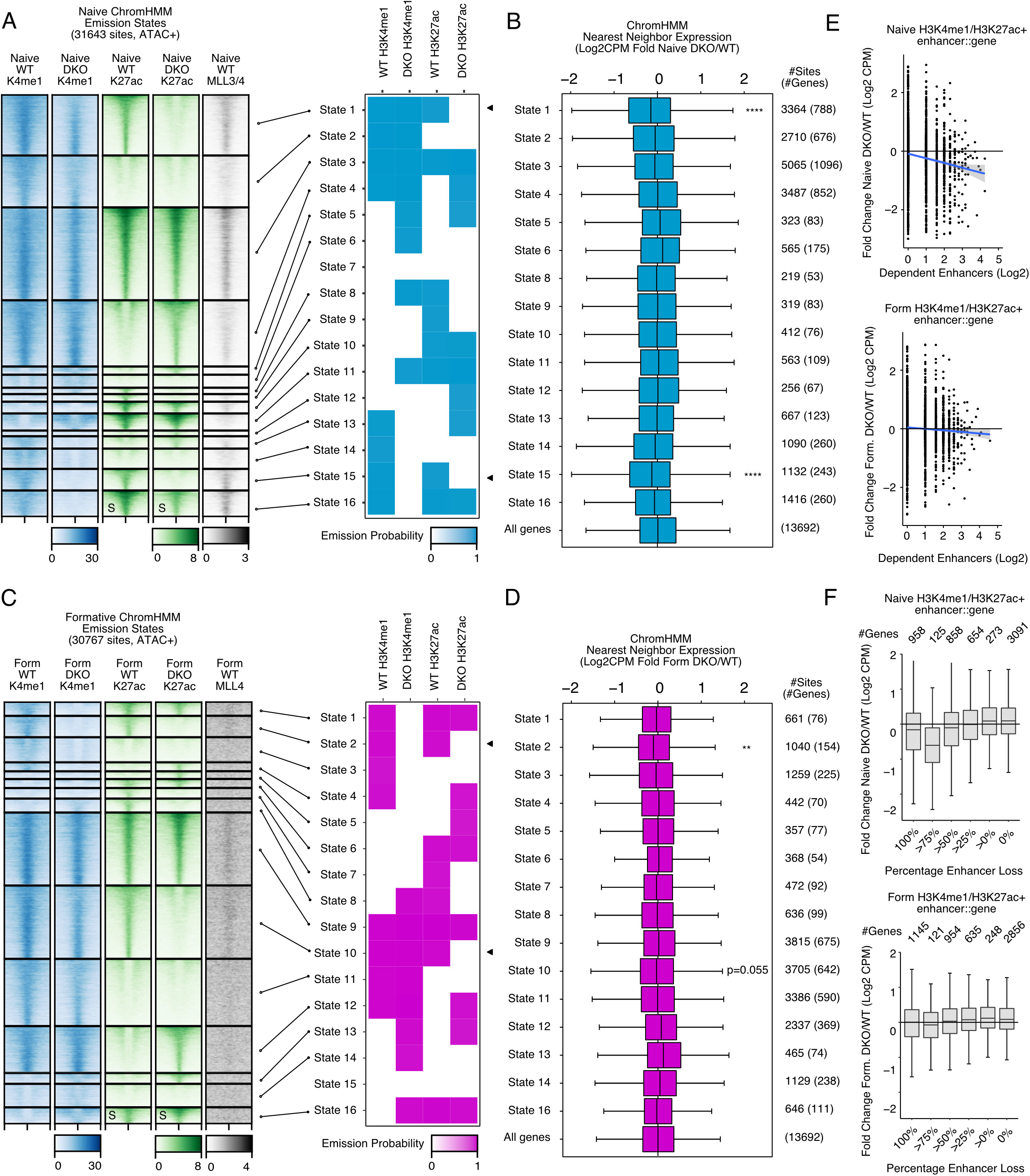
Distal H3K4me1 and H3K27ac are not functionally coupled with formative transcriptional activation. A) Naive ChromHMM emissions for 16 states yields all combinations of H3K4me1 and H3K27ac in WT and DKO. Heatmaps are clustered by each chromatin state. Black arrows denote states with MLL3/4 dependent H3K27ac. B) Gene expression changes of nearest TSS for all sites in each given state. Monte Carlo permutation sampling of “all genes control” to perform Mann-Whitney U Test for each state. Benjamini-Hochberg corrected. Selected statistics shown, additional statistics provided in supplemental table. C) Same as A with Formative samples. D) Same as B with Formative RNA-seq. All heatmap values and range are in CPM. For metagene analysis the range in CPM is the same as shown in heatmap for each factor. ChromHMM states without any H3K4me1 or H3K27ac emission probability are excluded. E) For all H3K4me1/H3K27ac+ sites in either Naive (top panel) or Formative (bottom panel), the total number of MLL3/4 dependent enhancers for a gene compared with the fold change of RNA levels DKO/WT in Log2CPM. Each dot represents one gene. Blue line represents generalized linear model, gray 95% confidence interval. F) Boxplots of relative expression DKO/WT of RNA levels for all genes associated with any H3K4me1/H3K27ac+ peak in either the naïve (top panel) or formative state (bottom panel). Each gene is binned by the percentage of their associated enhancer loss in DKOs. *p<0.05 **p<0.01 ***p<0.001 ****p<0.0001

Genes can be regulated by multiple enhancers(36, 37). Thus, we hypothesized that MLL3/4 independent enhancers may compensate for loss of MLL3/4 dependent enhancers explaining the underwhelming impact on expression. To test this hypothesis, we compared the change in expression (naive or formative DKO/WT) relative to the number of enhancers impacted by loss of MLL3/4 for each gene and their associated enhancers. In naive cells, this approach showed a correlation with the change in nearby gene expression (Fig.5E). In formative cells, only a very small effect was seen and only for genes with the greatest numbers of impacted enhancers. We obtained similar results by analyzing the proportion of enhancers that were lost per gene in each pluripotent state (Fig.5F). The contrast between naive and formative was even more striking when only considering sites within 10kb of the TSS (Fig.S6C,D). Therefore, compensation by non-MLL3/4 dependent enhancers does not appear to be a major factor rescuing gene expression in formative cells.

### Gene-centric analysis reveals a subset of distal loci associate with MLL3/4 dependent formative genes

Our data showed a requirement for MLL3/4 for H3K4 monomethylation at all sites that normally gain or lose the mark during the naive to formative transition (Fig. 2D,J,Fig.6A). They also showed a requirement for MLL3/4 for H3K27ac at 66% of sites that gain and 66% that lose acetylation during the transition (Fig3G,H, Fig.6A, Fig.S7A). However, only 13% of formative genes that normally increase in expression in the transition and only 39% naive genes that are normally downregulated were impacted by loss of MLL3/4. Among the naive genes that were impacted, there was a strong enrichment for cytokine pathways (Fig.S7B). Among the formative genes that were impacted, there was an enrichment for development related pathways including regionalization, migration, and morphogenesis (Fig.6B). This later finding suggested a role for initiating transcription of genes involved in a later stage of development, specifically gastrulation.

**Figure 6:**
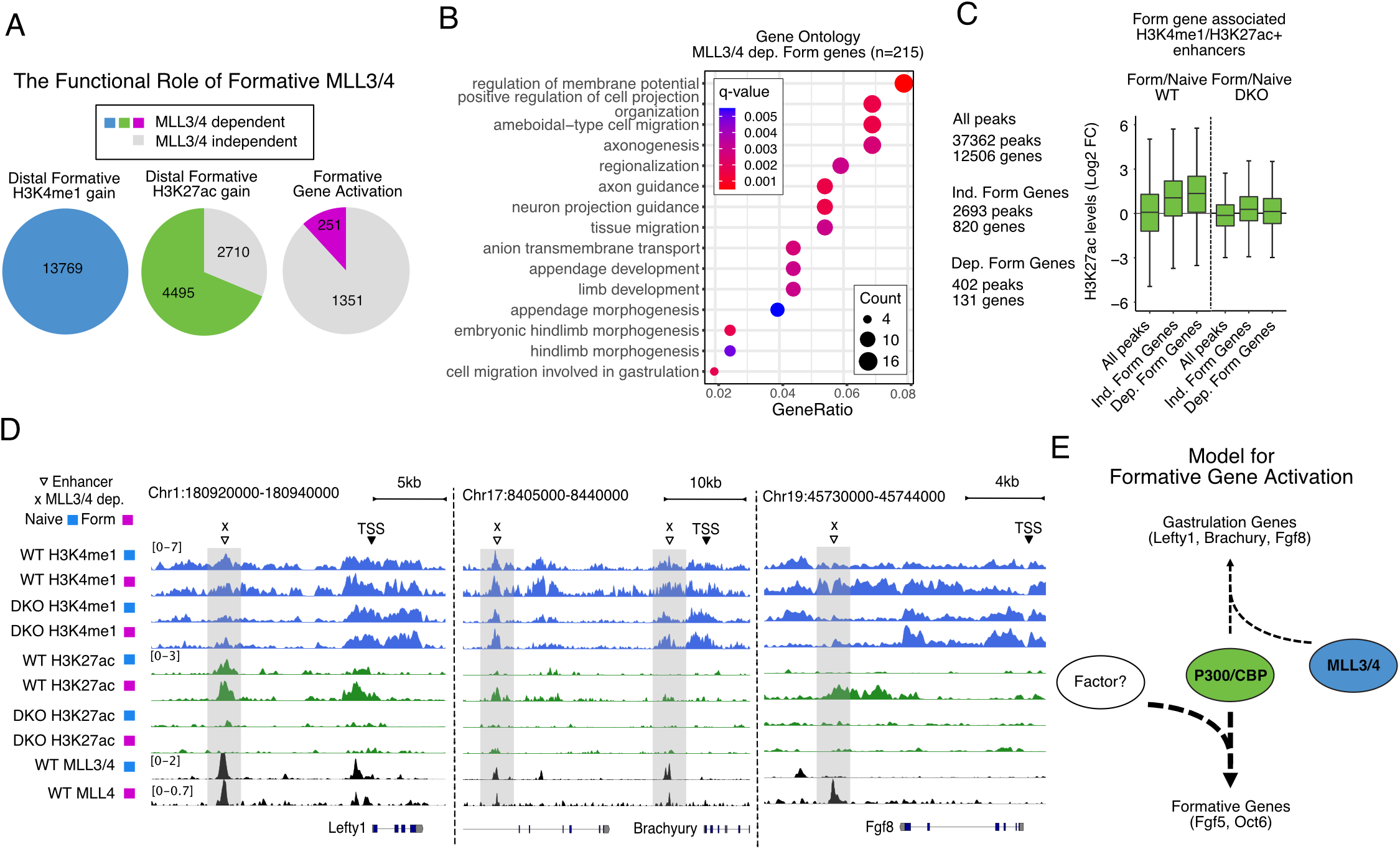
Gene-centric analysis reveals a subset of distal loci associate with MLL3/4 dependent formative genes. A) Pie charts showing fraction of normally gained H3K4me1, H3K27ac, or gene expression that are dependent on MLL3/4 or not. 215 formative genes represent those of the 283 formative down genes from Fig.1K without preexisting naive expression defects (Log2CPM naive DKO/WT > -1). B) Clusterprofile analysis of Biological Processes Gene Ontology for 215 MLL3/4 dependent formative genes. C) Fold change Log2CPM of H3K27ac density for H3K4me1/H3K27ac+ enhancers associated with formative genes that gained expression during transition in an either MLL3/4 independent or dependent fashion. Only 131 out of 215 dependent genes and 820 out of 1387 independent genes were able to be associated with an enhancer. D) Genome tracks for known and predicted formative targets of MLL3/4 Lefty1, Brachyury/T, and Fgf8. E) Model for the role of MLL3/4 independent and dependent mechanisms in the regulation of gained gene expression during the naïve to formative transition.

Our previous analyses utilized an enhancer-centric approach to identify changes in nearby genes. We next asked if impacted genes could be connected to nearby enhancers that showed loss of H3K27ac. To take this gene-centric approach, we categorized all genes that were normally upregulated during the transition to the formative state into those that were not impacted (independent) vs. impacted (dependent) by MLL3/4 loss. Among formative genes, this resulted in 820 independent formative genes and 131 dependent formative genes which could be linked to 2693 and 402 nearby H3K4me1/H3K27ac+ enhancers respectively. As a background control, we linked 12506 expressed genes from our RNA-seq to an associated 37362 H3K4me1/H3K27ac+ peaks. Both the dependent and independent formative genes normally showed an increase in H3K27ac at nearby enhancer sites during the naive to formative transition (Fig.6C). This increase in acetylation was mostly lost in DKO cells, whether or not the nearby genes failed to increase gene expression. Similarly, both dependent and independent naive genes (normally downregulated during transition) showed a decrease in nearby enhancer H3K27ac during the naive to transition. However, H3K27ac at enhancers nearby dependent naive genes was more impacted than those nearby independent genes in the DKO cells (Fig.S7C). Therefore, the gene-centric approach was consistent with enhancer-centric approach showing a more prevalent role of MLL3/4 regulated chromatin changes on the maintenance of naïve gene expression than the gain of formative gene expression.

To look deeper, we investigated the chromatin states nearby the small number of genes that were dependent on MLL3/4 for upregulation of transcript expression during the naive to formative transition by creating genome tracks (Fig.6D). These dependent genes included important gastrulation genes such as *T*/*Brachyury*, *Fgf8*, and *Lefty1*, the latter of which is a previously identified target of MLL3/4 (Wang et al. 2016). All three of these genes had one or more nearby enhancers that showed a loss of H3K4me1 and H3K27ac in MLL3/4 DKO formative cells. Thus, the expression of these key regulators relies on enhancers which require MLL3/4 for proper spatiotemporal regulation. However, they represent a very small subset of the thousands of enhancers that lose active marks upon the loss of MLL3/4. All together, these data show that from both a gene-centric or enhancer-centric perspective, H3K4me1 and/or H3K27ac at distal sites are largely uncoupled from target gene activation during the naïve to formative transition with the loss of MLL3/4.

## Discussion

Enhancers are thought to play a central role in gene regulation during cell fate transitions. Yet, the molecular epistasis of enhancer activation is incompletely understood. The prevailing model proposes that H3K4me1 deposited by MLL3/4 precedes H3K27ac deposition by P300/CBP and these events stimulate transcription. By combining state-of-the-art CUT&RUN/CUT&TAG technologies and knockouts of both MLL3 and MLL4, we evaluated this model during the well-defined transition from naive to formative pluripotency, where hundreds of genes are both silenced and activated. Surprisingly, the majority of these gene expression changes including de novo expression of many genes were unaffected by MLL3/4 loss. In contrast, loss of MLL3/4 led to large effects on the active histone modification signatures at thousands of enhancers. These results suggest that MLL3/4 and its orchestration of post-translational histone marks, play a relatively minor role in gene regulation during early ESC differentiation.

MLL3/4 are thought to be the major H3K4 monomethyltransferases in mammals(1, 2). However, we find that the majority of H3K4me1 is unaffected by MLL3/4 loss in both the naive and formative states. Consistent with our findings, similar results have been previously described in naive ESCs(12). By investigating the transition to the formative state, we were able to make several novel discoveries though. We discovered that all distal sites that normally show dynamic changes in H3K4me1, either gained or lost, were fully dependent on MLL3/4 for monomethylation. In contrast, only a small fraction of the sites that normally do not change during the transition showed any requirement for enzymes in maintaining monomethylation levels. Therefore, there appear to be at least two distinct types of cis-regulatory elements with regard to MLL3/4 dependency. This raises important follow-up questions such as what enzymes are responsible for the predominant group of H3K4me1 sites that do not require MLL3/4 for maintenance? MLL2 has been suggested as an alternative monomethyltransferase in naive ESCs, but only at a small number of sites that co-exist with H3K27m3, a repressive mark(12, 38). MLL1 represents another potential candidate. Interestingly, MLL1 has been shown to inhibit the reprogramming of primed pluripotent (EpiSCs) to naive ESCs(39). However, its role in regulating H3K4me1 in naive cells during differentiation has not been evaluated. Other candidates include SMYD1 and SMYD2 which are SET domain containing proteins that have previously been shown to be required for H3K4me1 and or H3K4m2 at de novo enhancers during macrophage activation(40). Additionally, whether there are distinct roles for the MLL3/4 dependent vs. independent distal sites remains unclear. Given most enhancer activation in the absence of MLL3/4 occurs at these independent H3K4me1 sites they may represent a collection of primed sites distinct from MLL3/4. Surprisingly though, previous studies have shown that at least in the case of MLL3/4, the proteins, but not their enzymatic activity are important in CBP/P300 recruitment and H3K27ac(12). Therefore, it will be important to understand the link between the factors resulting in methylation of these MLL3/4 independent sites and those that deposit H3K27ac at these sites.

MLL3/4 are thought to be important recruiters of CBP/P300, whose catalytic byproduct H3K27ac is a key marker of enhancer activation. In our data, we identify that more than half of H3K27ac distal sites lose acetylation with MLL3/4 loss, with a greater proportion at dynamic sites than unchanging sites. Many sites that lose H3K27ac do so independent of H3K4me1 loss. Whether this represents an indirect role or a direct role for MLL3/4 in the recruitment of CBP/P300 at sites that also retain H3K4me1 is unclear. However, the presence of MLL3/4 binding at many of these sites supports a direct role. In addition to the MLL3/4 dependent sites, there is also a large number of MLL3/4 independent H3K27ac sites, including many occurring at regions that do require MLL3/4 for H3K4me1. This indicates multiple mechanisms for CBP/P300 recruitment to enhancers that are differentially used depending on the specific enhancer and chromatin state. How different enhancers recruit CBP/P300 using distinct mechanisms is unclear but is likely in part due to the underlying transcription factors binding to those enhancers. Enhancer subcategories consisting of varying types of TFs have recently been shown to require specific chromatin complexes for enhancer activity(41). Consistent with the role for different TFs underlying these requirements, TF motif enrichment analysis uncovered several TFs whose motifs were highly enriched at each distinct category of MLL3/4 dependent and independent enhancers. CBP/P300 could either be directly recruited by these TFs and/or be recruited through other factors compensating for loss of MLL3/4. Future studies are necessary to fully elucidate the requirements for recruitment of specific epigenetic complexes by distinct TFs.

Ultimately, it is thought that enhancer activation as defined by the gain of H3K27ac drives increased expression of its target gene. Surprisingly though, while we observe major impacts on the chromatin state of sites with dynamic H3K4me1 and H3K27ac during the naive to formative transition, there are relatively modest changes in the gain and loss of gene expression. The activation of the formative transcriptional program is largely unaffected, with only a few exceptions such as Lefty1 and Fgf8. The key markers of the formative state including Otx2, Oct6, and Fgf5 are all normally upregulated. This unexpected finding raised the question whether there was any specific MLL3/4 dependent enhancer phenotype in formative cells that could be linked to a failure to activate gene expression. Analysis of genes either nearby MLL3/4 dependent formative H3K4me1 sites or MLL3/4 dependent formative H3K27ac sites during the transition did not identify a reduction in gene activation. Most surprising, interrogation of genes nearby enhancers that fail to gain both H3K4me1 and H3K27ac also showed normal increases in gene expression. Only when considering the small number of sites where both H3K4me1 and H3K27ac were depleted, independent of their dynamics, was there a small but significant effect for a handful of genes. Consistent with our data, recent work mutating H3 and/or H3.3 Lysine 27 sites to Arginine, which cannot be acetylated did not impact either naive gene expression or the activation of the formative program in differentiating ESCs(14, 15). However, these findings are interpreted as a lack of a requirement for H3K27ac rather than an inability to recruit enhancer CBP/P300 at large where we also expect a loss in other active enhancer histone acetylations deposited by CBP/P300 such as H2B acetylation(42). Either way, it is unclear whether many of the sites dependent on MLL3/4 for active histone marks are dispensable for gene expression changes and or these sites are functional enhancers without an active histone signature. Of note, loss of MLL3/4 can also impact the recruitment of chromatin remodelers (i.e. the BAF complex) and enhancer-promoter looping, making the minimal impact MLL3/4 loss has on gene transcription during this transition even more surprising (9,43,44).

MLL4 KO embryos die shortly following gastrulation but show defects starting in early gastrulation(6, 21). In contrast, MLL3 KO embryos die perinatally due to developmental defects in the lung. MLL3/4 double KO embryos phenocopy the MLL4 KO consistent with the MLL4 being able to compensate for MLL3 loss in early development. The direct molecular basis for these phenotypes is poorly understood. However, it is important to note that even though MLL3/4 loss had little impact on gene activation during the ESC transition from naive to formative pluripotency, many genes were dysregulated in steady state naive ESCs and remained so in the formative state. In the case of persisting naive genes (Fig. 1K), this is consistent with previous reports that MLL4 KOs fail to repress naive genes in LIF cultured cells after withdrawal of 2i from naive (LIF+2i) culture conditions(45). Some misregulated genes are likely direct targets given the downregulation of genes nearby MLL3/4 dependent enhancers upon loss of the enzymes. In addition, there are likely indirect effects potentially including additional unknown non-transcription related functions for MLL3/4. It is also important to note that other studies have shown important roles for MLL3/4 in the upregulation of gastrulation markers in the setting of embryoid body differentiation, a model for a later stage in early development than studied here(22). Even later in development, MLL3/4 is essential for developmental programs in a variety of tissues including cardiac, neural crest, muscle, and adipose specific programs(6,7,46,47). The differences between these studies and ours suggest a cell-context specific function of MLL3/4. A potential reason is the compendium of TFs that require or do not require MLL3/4 is different for each transition. For instance, PPAR*γ* and CEBP*α* are major adipogenic factors that require MLL3/4 to function, while at least some of the pluripotency factors and the drivers of formative state are MLL3/4 independent based on our data. One possibility may be that transcriptional roles of MLL3/4 proteins are limited to determinants of differentiation pathways post-pluripotency and is most important in specifying somatic cell fates. Interestingly, a recent preprint suggests that MLL3/4 enzymatic activity is also important for a subset of cell fates such as extraembryonic endoderm and trophectoderm but not germ layer formation(48). Taken together, these data suggest fundamental and currently underappreciated differences in transcriptional control between pluripotent and somatic cell types.

## Conclusions

In sum, our data demonstrates that cell-type specific enhancer activation is not uniquely performed by MLL3 and MLL4. Instead, gene regulation appears to be partitioned during the naive to formative transition. MLL3/4 is required for a small subset of genes critical for gastrulation and later developmental processes, but formative transcriptional activation is mediated by other mechanisms including MLL3/4 independent enhancers (Fig.6E). By identifying rules and exceptions to the current model of enhancer activation we highlight current gaps in knowledge concerning the molecular steps regulating these processes. Further investigation into the molecular interplay and steps governing enhancer function will advance our understanding of gene regulation in health and gene dysregulation in disease.

## Methods

### Cell Culture

Mouse ESCs were cultured in Knockout DMEM (Thermo Fisher, CAT#10829018) supplemented with 15% FBS, L-Glutamine, Penicillin/Streptamycin, NEAA, LIF (1000U/mL), and 2i (1uM MEK inhibitor PD0325901 and 3uM GSK3 inhibitor CHIR99021). Wildtype and dCD cells were R1 ESCs from a 129 strain background. MLL3^-/-^; MLL4^fl/fl^ ESCs were a mixed background of C57BL/6J and 129 strains. Formative cells were generated by removal of LIF and 2i. Briefly, 5e4 ESCs were plated per well of a 6 well plate on day −1 in LIF+2i media. To initiate differentiation, LIF and 2i were removed 24 hours after seeding (Day 0). Formative cells were collected on day 3 of differentiation, 63 hours after removal of LIF and 2i. To overcome proliferation defects 7.5e4 MLL3/4 DKO cells were plated per well of a 6 well. Naive cells were passaged and staged appropriately for simultaneous harvest. The expected amino acid substitutions in MLL3/4 dCD cells were validated using Sanger sequencing on amplicons generated from genomic PCR. To generate DKOs from MLL3^-/-^; MLL4^fl/fl^ ESCs we transfected with Rosa26-CRE-ERT2 plasmid linearized by EcoRI, selected with puromycin, and genotyped using genomic PCR. We then added tamoxifen for 3 days and picked clones to validate knockout of MLL4 floxed allele. Primers for genotyping listed in the supplemental table. Lines consistently tested negative for mycoplasm.

### Crystal Violet Assay

To assess proliferation cells were plated in 24 well plates at approximately 12.5e4 cells per well. After 24, 48, and 72 hours, cells were then washed with PBS and 200ul Crystal Violet (0.2% Crystal Violet, 2% ethanol in dH2O) was added to the plate for 10 minutes at room temperature. Cells were then washed twice by gently submerging plate in tap water before 600ul 1% SDS was added to the well to solubilize the stain. Plate was placed on a shaker until color was uniform in the well and 200ul was transferred to a 96 well plate for reading absorbance at 570nm with a plate reader (SpectraMax M5). Absorbance for each time point was normalized to a blank well.

### qPCR and analysis

To perform qPCR we first extracted RNA by adding Trizol directly to plates. After adding chloroform, an isopropanol precipitation with GlycoBlue was performed followed by ethanol washes. RNA pellets were resuspended in RNAse-free water and quantified using a NanoDrop. 200ug of RNA was used for cDNA synthesis using Maxima First Strand Synthesis Kit (Thermo Fisher, CAT#K1672) with half reactions according to the manufacturer’s protocol. qPCR on cDNA was performed using SYBRgreen master mix (Applied Biosystems, CAT#A25742) using 6ul final volume on a QuantStudio5 qPCR machine (Applied Biosystems). qPCR primers for targets are listed in the supplemental table. Target Ct values were normalized to GAPDH internally for each sample and then set relative to the naive WT negative control.

### Protein extraction and Westerns

To harvest protein for western assays, cells were trypsinized, washed once with ice cold PBS before adding RIPA buffer with protease inhibitors (Sigma CAT#P8340). After 15 minutes on ice, lysed cells were centrifuged at 16000g for 10 minutes and the supernatants representing whole cell fractions were collected and snap frozen in liquid nitrogen. Protein quantification was conducted using a Micro BCA protein assay kit (Thermo CAT#23235). 40ug of protein was loaded per well of SDS PAGE gels for western. For histone purification we utilized the acid extraction protocol exactly as described in Shechter et al. 2007 using 5e6 cells as input(49).

Westerns were typically conducted using a Bio-Rad system with Tris-Glycine gels purchased from Bio-Rad and transferred to methanol activated PVDF membranes. Membranes were blocked, and stained with primaries and secondaries using Li-Cor Odyssey Blocking Buffer (Li-Cor, CAT#927-60001) mixed 1:1 with TBS. Primary antibodies for western were Rabbit anti-MLL3 (provided by Kai Ge), Rabbit anti-MLL4 (provided by Kai Ge), Mouse anti-OCT4 (BD Biosciences, CAT#611202), Rabbit anti-NANOG (Cell Signaling, CAT#61419), Rabbit anti-H3K4me1 (Abcam, CAT#ab8895), and Mouse anti-H3 (Cell Signaling, CAT#3738). Westerns for histones were conducted similarly to other targets except gels were transferred to PVDF membranes using CAPS buffer (500mM CAPS, adjusted to pH 10.5 with NaOH).

High molecular weight westerns for MLL3 and MLL4 were conducted using a NuPage SDS-PAGE and Transfer system with an XCell electrophoresis unit according to manufacturer’s protocols with several modifications (Thermo Fisher, CAT#E10002). In brief, 40ug of whole cell protein lysate was incubated with LDS loading buffer (Thermo Fisher, CAT#NP0007, added BME to 1% final concentration) and incubated at 70C for 10min. Samples were loaded into 3-8% Tris-acetate gels (Thermo Fisher, CAT#EA03752BOX) and ran in sample running buffer (Thermo Fisher, CAT#LA0041) supplemented with NuPage Antioxidant (Thermo Fisher, CAT#NP0005). Samples were run at 80V for 30min and then 120V for 120min. Samples were then transferred to PVDF membranes using NuPage Transfer Buffer (Thermo Fisher, CAT#NP0006, with added 10% Methanol, NuPage Antioxidant, 0.01% SDS) at 30V in a cold room for either 2 hours or overnight.

### RNA sequencing

Total RNA was extracted and purified from cells using Trizol followed by ethanol precipitation. RNA-seq libraries were generated using the QuantSeq 3’ mRNA-Seq Library Prep Kit FWD for Illumina (Lexogen, CAT#A01172) according to their protocol using 200ng of total RNA for input. We utilized the PCR Add-on kit for Illumina (Lexogen, CAT#M02096) to determine an appropriate number of PCR cycles to amplify libraries. Amplified libraries were quantified using Agilent Tapestation 4200. Libraries were pooled and sequenced using a HiSeq 4000 to obtain single end 50bp reads. At least 10 million mapped reads or more per sample were obtained.

### RNA sequencing processing and analysis

Adapters for sample FastQ files were trimmed using cutadapt v2.5 followed by alignment to the mm10 genome using STAR align 2.7.2b. Featurecounts from Subread v1.6.4 was used to generate a counts matrix of reads per gene. FastQC v0.11.8 and MultiQC were used to validate quality sequencing and mapping.

The gene count matrix was converted to CPM and filtered for genes greater than 1 cpm in at least two total samples. Samples were normalized using TMM. Next, Log2 CPM averages were calculated for replicates of each sample for scatterplot visualization and nearest neighbor TSS analysis. All transcriptomics analyses were conducted using CPM values from TMM normalization of all samples except for nearest neighbor analysis where WT and DKO were TMM normalized together. To conduct differential gene expression we performed DESeq2 v1.34.0 analysis using the raw gene count matrix as input for each desired comparison. Gene ontology was performed using ClusterProfiler 4.2.2. Custom R code for other downstream transcriptomics analyses and visualization provided on Github.

### CUT&RUN sequencing and processing

CUT&RUN was conducted using the protocol from Skene et al. 2018 with the following modifications: Freshly trypsinized cells were bound to activated Concanavalin A beads (Bang Laboratories, #BP531) at a ratio of 2e5 cells/10ul beads in CR Wash buffer (20mM HEPES, 150mM NaCl, 0.5mM Spermidine with protease inhibitors added) at room temp. An input of 2e5 cells were used per target. Bead-bound cells were then incubated rotating overnight at 4C in CR Antibody buffer (CR Wash with 0.05% Digitonin, 2mM EDTA) containing primary antibody. We used the following antibodies for CUT&RUN: 1:100 Rabbit anti-H3K4me1 (Abcam, CAT#ab8895), Rabbit anti-MLL4 1:100 (provided by Kai Ge), and 1:100 Rabbit IgG isotype control (Abcam, CAT#ab171870). After primary, we washed 3 times 5 minutes each with cold CR Dig-wash buffer (CR Wash with 0.05% Digitonin) and incubated with pA-MNase (1:100 of 143ug/mL provided by Steve Henikoff) for 1 hour rotating at 4C. After MNase binding, we washed 3 times 5 each with cold CR Dig-wash buffer, and chilled cells down to 0C using a metal tube rack partially submerged in an ice water slurry. MNase digestion was induced by adding CaCl2 at a final concentration of 2mM. After 30 minutes of digestion, the reaction was quenched using Stop Buffer containing 340mM NaCl, 20mM EDTA, 4mM EGTA, 0.05% Digitonin, 100ug/mL RNAse A, 50ug/mL Glycogen, and approximately 2pg/mL Yeast spike-in DNA (provided by Steve Henikoff). The digested fragments for each sample were then extracted using a phenol chloroform extraction. Library preparation on samples was conducted using manufacturer’s protocols for NEBNext Ultra II Dna Library Prep Kit (New England BioLabs, CAT#E7645) and NEB Multiplex Dual Index oligos (New England BioLabs, CAT#E7600, #E7780) with the following modifications. We input approximately 10ng of sample for half reactions, we diluted the NEBNext Illumina adaptor 1:25, we used the following PCR cycling conditions: 1 cycle of Initial Denaturation at 98C for 30 seconds, 12+ cycles of Denaturation at 98C for 10 seconds then Annealing/Extension at 65C for 10 seconds, and 1 final cycle of extension at 65C for 5 minutes. Following library preparation, double size selection was performed using Ampure beads and quality and concentration of libraries were determined by an Agilent 4200 Tapestation with High-Sensitivity D1000 reagents before pooling for sequencing.

Fastq files for CUT&RUN samples were processed using Nextflow(50) and the nf-core CUT&RUN pipeline v1.0.0 beta. In brief, adapters were trimmed using Trim Galore. Paired-end alignment was performed using Bowtie2 and peaks were called using SEACR with a peak threshold of 0.05 using spike in calibration performed using the E.coli genome K12. However, all downstream analysis was performed using CPM normalized samples. To reduce high background observed with MLL4 antibody we subtracted MLL4 CUT&RUN in DKO cells away from WT to generate formative MLL4 CUT&RUN bigWigs for use in heatmaps.

### CUT&TAG sequencing and processing

CUT&TAG was conducted using the protocol from Kaya-Okur et al. 2020 with the following modifications: freshly trypsinized cells were bound to Concanavalin A beads at a ratio of 2e5 cells/7ul beads in CR wash at room temp. We used 2e5 cells as input per sample. Bead-bound cells were then incubated rotating overnight at 4C in CT Antibody buffer (CR Wash with 0.05% Digitonin, 2mM EDTA, 1mg/mL BSA) containing primary antibody. We used the following primary antibodies for CUT&TAG: 1:100 Rabbit anti-H3K27ac (Abcam, ab4729) and 1:100 Rabbit IgG isotype control (Abcam, ab171870). After primary, samples were washed 3 times for 5 minutes each using CR Dig-wash buffer and resuspended in 1:100 secondary antibody (Guinea pig anti-rabbit, Antibodies Online #ABIN101961) in CR Dig-wash buffer at 4C for 1 hour rotating at 4C. Samples were then incubated for 1 hour at 4C with 50ul of approximately 25nM homemade pA-Tn5 in CT Dig300 wash buffer (20mM HEPES, 300mM NaCl, 0.01% Digitonin, 0.5mM Spermidine with Roche cOmplete protease inhibitors added). Recombinant Tn5 was purified and loaded with adapters as previously described (Kaya-Okur et al. 2019). After Tn5 incubation, samples were washed 3 times for 5 minutes each with CT Dig300 wash buffer. Tagmentation was then initiated for 1hr at 37C in a thermocycler by adding MgCl2 to 10mM final concentration in 50uL volume. The tagmentation reaction was quenched immediately afterwards by adding 1.6ul of 0.5M EDTA, 1ul of 10mg/mL Proteinase K, and 1ul of 5% SDS. Samples were then incubated at 55C for 2 hours in a thermocycler to denature Tn5 and solubilize tagmented chromatin. After incubation, samples were magnetized and the supernatant was transferred to new wells where SPRI bead purification was performed using homemade beads to select all DNA fragment lengths larger than 100bp. Samples were eluted in 0.1X TE and approximately half of each sample was used for library preparation using NEBNext HIFI Polymerase with custom indices synthesized by IDT. An appropriate number of cycles for each target was chosen to prevent overamplification bias. After amplification, libraries were purified with 1.2x homemade SPRI beads to select for fragments >250bp and eluted in 0.1X TE. Quality and concentration of libraries were determined by an Agilent 4200 Tapestation with D1000 reagents before pooling for sequencing.

CUT&TAG samples were processed similarly to CUT&RUN samples using the same Nextflow pipeline.

### ATAC-seq sequencing and processing

ATAC-seq libraries were generated using the Active Motif commercial kit following the provided protocol with kit components (Active Motif, CAT#53150): Specifically, freshly trypsinized cells were washed with ice cold PBS using 75K cells as input for each sample. Cells were lysed using ice cold ATAC Lysis Buffer and we added 50ul of Tagmentation Master Mix to each sample. We incubated tagmentation reactions at 37C for 30min in a thermomixer set to 800 rpm. We immediately transferred samples to a new tube and performed DNA column purification. After purification we amplified the libraries using the following PCR conditions: 1 cycle at 72C for 5 minutes, 1 cycle at 98C for 30 seconds, and finally 10 cycles of 98C for 10 seconds, 63C for 30 seconds, 72C for 1 minute. Amplified libraries were purified using 1.2x SPRI bead solution. Quality and concentration of libraries were determined by an Agilent 4200 Tapestation with D1000 reagents before pooling for sequencing.

Fastq files for ATAC-seq samples were processed using Nextflow and the nf-core ATAC-seq pipeline v1.2.1. In brief, adapters were trimmed using Trim Galore. Paired-end alignment was performed using BWA and peaks were called using MACS2 in broad peak mode with a cutoff value of 0.1. BAM files were converted to bigWigs. Using read counts from consensus peaks of WT and DKO samples we performed TMM normalization using EdgeR to generate scale factors used during bigWig generation of samples.

### ChIP-seq processing

Fastq files for ChIP-seq from previously published studies were processed using Nextflow and the nf-core ChIP-seq pipeline v1.2.1. In brief, adapters were trimmed using Trim Galore. Paired-end alignment was performed using BWA, duplicates were removed, and peaks were called using MACS2 in broad peak mode with a cutoff value of 0.1. We subtracted the MLL3/4 ChIP-seq in DKO cells away from WT samples to generate naive MLL3/4 ChIP-seq bigWigs for use in heatmaps.

### Peak analysis

We performed differential signal enrichment analysis using Diffbind on SEACR peaks derived from CUT&RUN or CUT&TAG samples. We then used the output from Diffbind, containing the union of input SEACR peaks, for two packages: 1) We used Deeptools multiBigwigsummary to count reads at each peak for the different targets/samples and 2) we used HOMER annotatePeaks to identify nearest gene and annotate genomic features. These files were then imported into R and joined together by peak coordinates to make a dataframe which we used to filter for phenotypes and calculate fold changes. After genomic feature annotation by Homer we classify intergenic and intronic peaks as “Distal”, TSS as “Promoter”, and everything else as “Other” including: exon, transcriptional termination sites (TTS), UTR regions, and those without assigned features. To calculate Log2CPM peak densities we took counts for each peak from multibigwigsummary multiplied them by a constant (1e6), added 1, and then performed Log2(x). Finally, we divided by the width in basepairs of the peak. Fold changes were conducted using these Log2CPM peak density values. H3K27ac CUT&TAG data was subset by H3K27ac peaks that overlapped at least 75% of an WT ATAC-seq peak using bedops. We also used bedops on ATAC subset H3K27ac sites to identify H3K27ac peaks that had overlapping H3K4me1 peaks to generate our H3K4me1/H3K27ac+ peak lists. ChromHMM segmentations were similarly filtered for chromatin states that overlapped at least 75% of an WT ATAC-seq peak.

## Declarations

### Ethics approval and consent to participate

Not applicable

### Consent for publication

Not applicable

### Availability of data and materials

The data generated and their analysis in the current study are available in the GEO Database using the identifier GSExxxxxx. Code generated for analysis will be hosted on Github or otherwise available by request.

### Author’s Contributions

Conceptualization: RMB, RB; Methodology: RMB; Validation: RMB; Formal Analysis: RMB; Investigation: RMB, KC; Resources: RB; Data Curation: RMB; Writing - original draft: RMB; Writing - review and editing: RMB, RB; Visualization: RMB; Supervision: RMB, RB; Project Administration: RB; Funding Acquisition: RB.

## Acknowledgements

We thank Kai Ge for providing the MLL3KO;MLL4 floxed ESC line and antibodies for MLL3 and MLL4. We thank Joanna Wysocka for providing the MLL3/4 catalytic dead cell line. We thank Steve Henikoff for sending purified pA-MNase. We would like to thank the Nextflow community for helpful advice on processing of sequencing samples including Chris Cheshire and Charlotte West for providing access to a beta for nf-core CUT&RUN pipeline. We would like to thank Bryan Marsh, Deniz Gökbuget, and Brian Deveale for helpful discussions throughout the project including proofreading the original manuscript. We would like to thank Dan Lim, Yin Shen, and Elphege Nora for their thoughtful comments and constructive feedback on the manuscript.

## Supplementary Figure Legends

**Figure S1:**
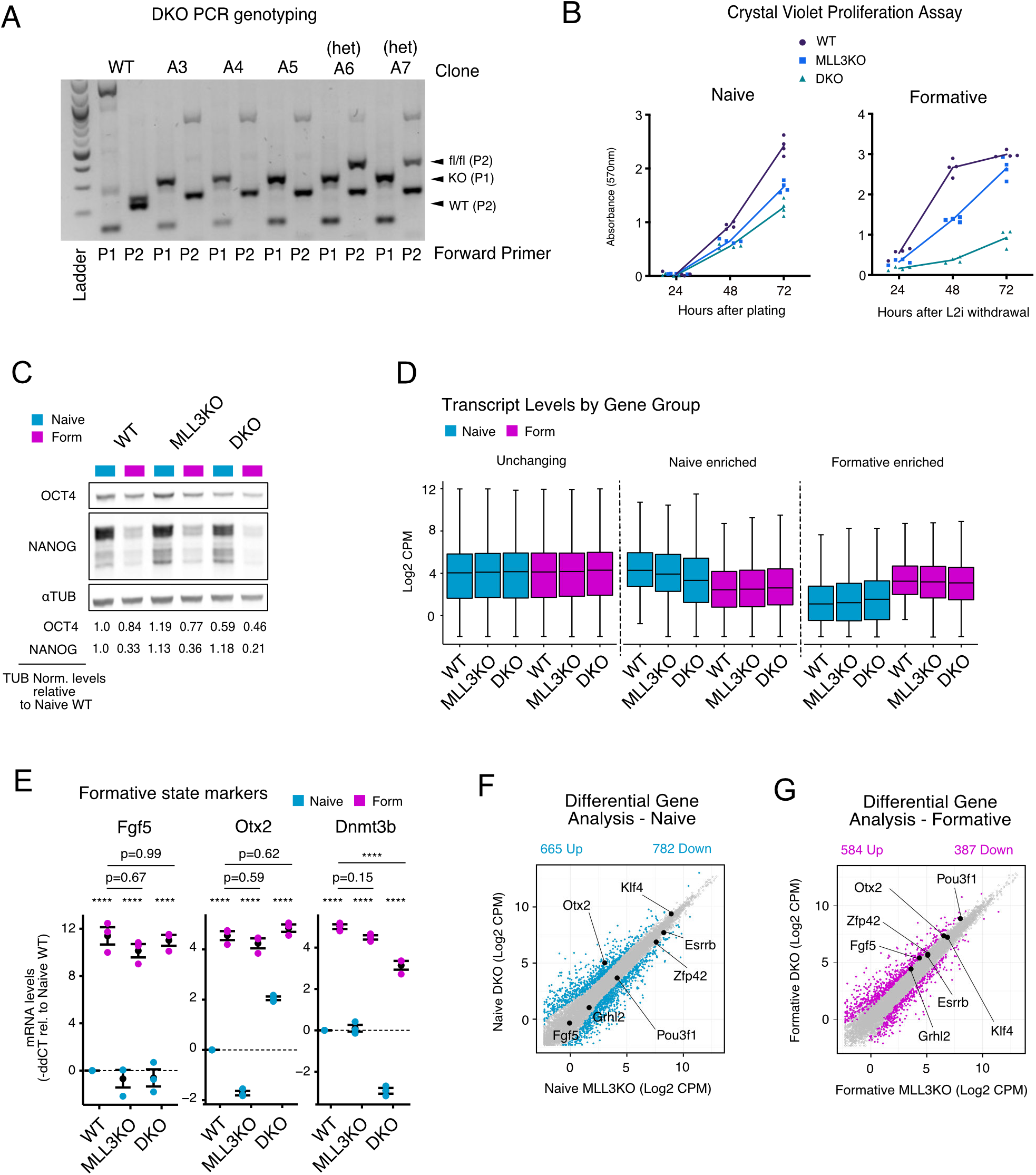
MLL3/4 is dispensable for transcriptional activation of much of the formative program. A) Confirmation of multiple DKO clones using genomic PCR following tamoxifen treatment. B) Proliferation measurements using crystal violet assays. C) Westerns on whole cell protein fractions of OCT4, NANOG and *α*TUBULIN loading control. Quantifications below are fold changes relative to naïve WT after normalizing OCT4 or NANOG for TUBULIN levels in each lane. D) qPCR of formative enriched markers Fgf5, Otx2, Dnmt3b. For each marker selected comparisons are shown resulting from a Two-way ANOVA, Tukey’s multiple comparison test for all samples. Mean and S.E.M. in black. Additional statistics provided in supplemental table. E) Boxplots of transcript levels in Log2CPM for unchanging, naïve enriched or formative enriched genes. F) DGE analysis on MLL3KO naive and DKO naive samples (significant genes colored, P.adj < 0.05 and Log2FC > 1). G) Same as F but comparing the formative state. *p<0.05 **p<0.01 ***p<0.001 ****p<0.0001

**Figure S2:**
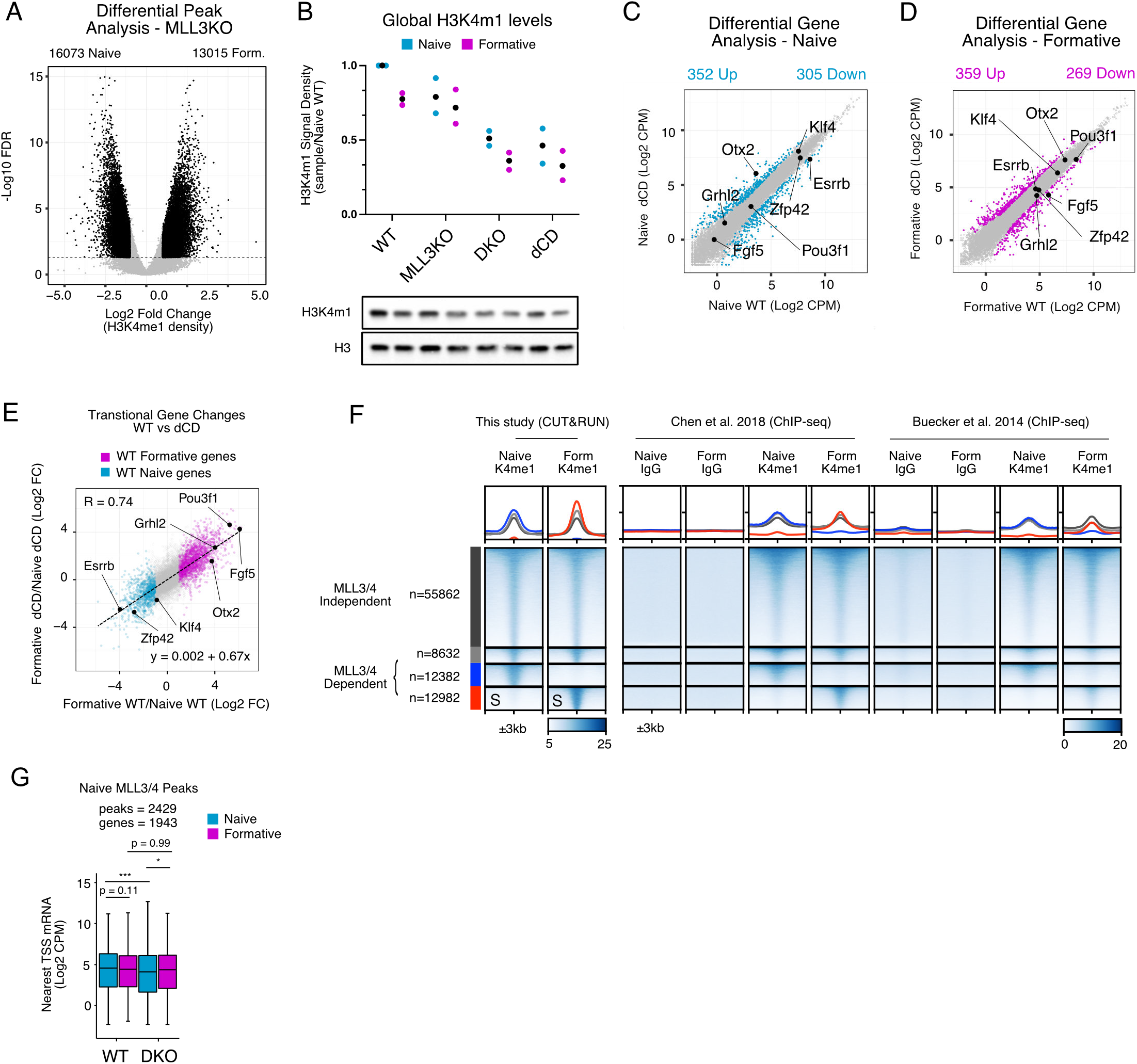
MLL3/4 is required for all dynamic H3K4me1 deposition during pluripotent transition. A) Diffbind analysis of H3K4me1 signal in MLL3KO cells at WT peaks. B) Quantifications of westerns on histones purified by acid-extraction. H3K4me1 levels are relative to naïve WT levels after first normalizing by H3 signal in each lane. Black dot represents mean (n=2). C) mRNA Log2CPM of naive MLL3/4 dCD cells vs. WT cells. Significant genes highlighted. D) same as B with formative samples. E) Foldchange (formative/naive) of all genes for MLL3/4 dCD compared to WT. WT formative and naive genes from WT DGE analysis colored. R, Pearson’s coefficient. Dashed line and linear equation represent linear model of all genes. F) Heatmap of ChIP-seq for H3K4me1 in naive and formative cells from two published studies. All heatmap values and range are in CPM. For metagene analysis the range in CPM is the same as shown in heatmap for each factor. G) Nearest neighbor TSS analysis of expression levels in Log2CPM for each RNA-seq dataset near naive MLL3/4 ChIP-seq peak categories. Multi-comparison paired Wilcoxon Rank-Sum Test, Benjamini-Hochberg corrected. *p<0.05 **p<0.01 ***p<0.001 ****p<0.0001

**Figure S3:**
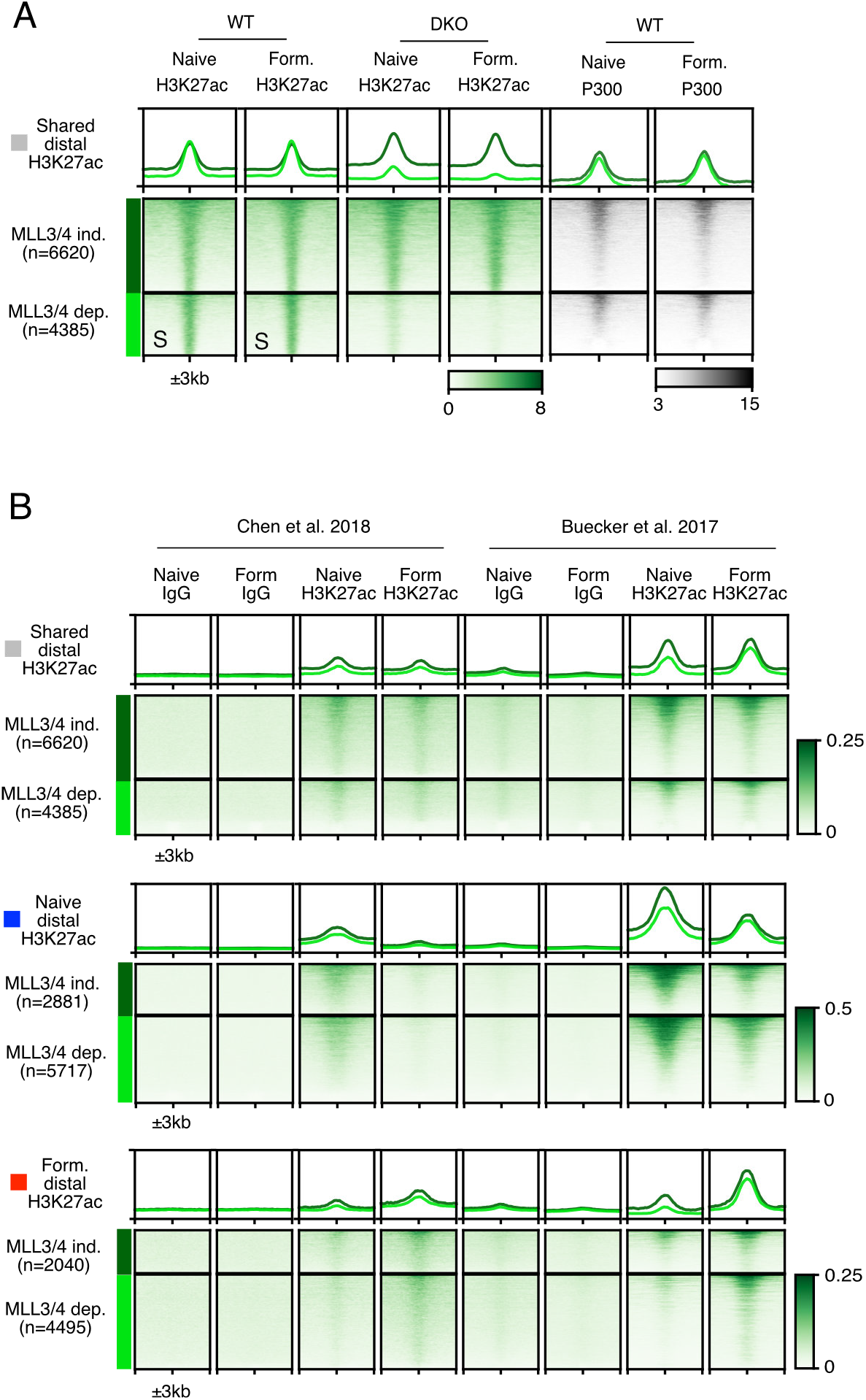
MLL3/4 dependent and independent distal H3K27ac deposition. A) Heatmap of shared H3K27ac sites clustered by MLL3/4 independent or dependent H3K27ac. B) Heatmaps of ChIP-seq for H3K27ac in naive and formative cells from two published studies. All heatmap values and range are in CPM. For metagene analysis the range in CPM is the same as shown in heatmap for each factor.

**Figure S4:**
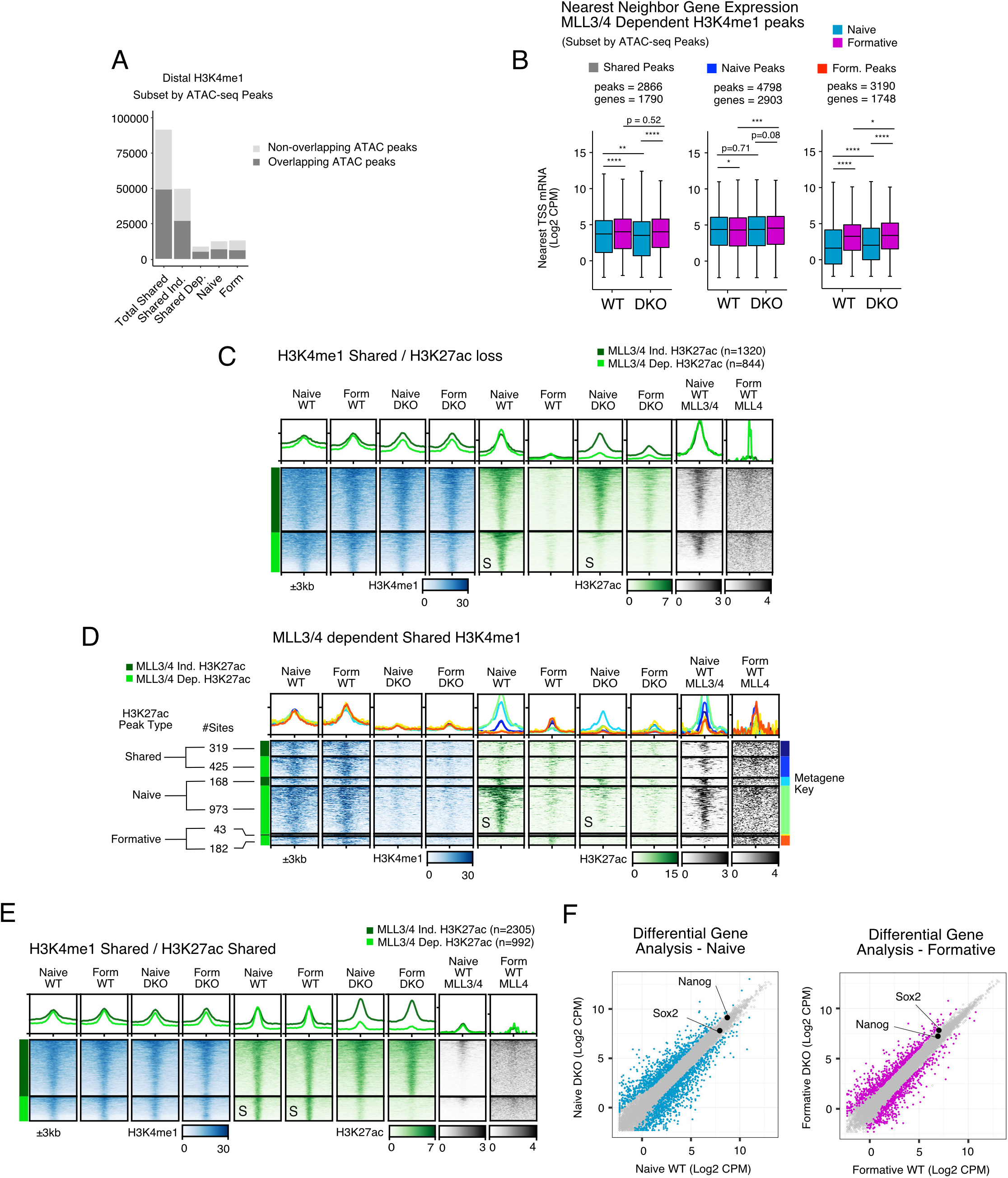
Enhancer Activation Can Occur Independently of MLL3/4. A) Peak count of H3K4me1 categories from Fig2. after filtering for distal H3K4me1 overlapping WT ATAC peaks. B) Nearest neighbor TSS analysis of expression levels in Log2CPM for each RNA-seq dataset near H3K4me1 peak categories after subsetting for ATAC peak overlap. Multi-comparison paired Wilcoxon Rank-Sum Test, Benjamini-Hochberg corrected. C) Heatmaps for shared H3K4me1 that overlaps with naive enriched H3K27ac, clustered by MLL3/4 independent or dependent H3K27ac. D) Heatmaps for MLL3/4 dependent H3K4me1 overlapped with sites that have shared, naive, or formative H3K27ac. Clustered additionally by MLL3/4 independent or dependent H3K27ac. E) Heatmaps for shared H3K4me1 that overlaps with shared H3K27ac, clustered by MLL3/4 independent or dependent H3K27ac. All heatmap values and range are in CPM. For metagene analysis the range in CPM is the same as shown in heatmap for each factor. F) Log2CPM of Sox2 and Nanog in naive and formative samples comparing WT and DKO cells. Sox2 is significantly up in the formative state. *p<0.05 **p<0.01 ***p<0.001 ****p<0.0001

**Figure S5:**
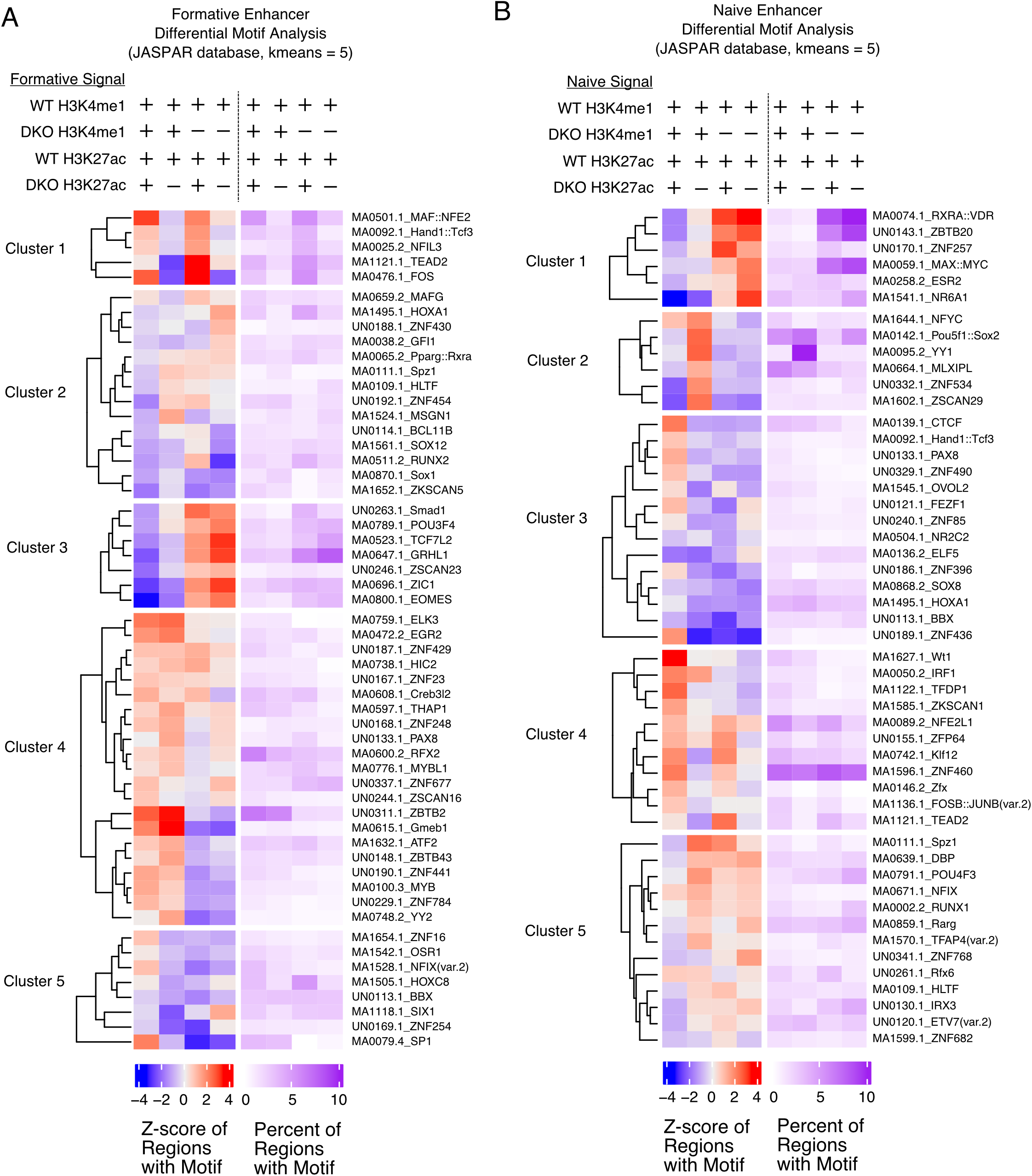
Enhancer Activation Can Occur Independently of MLL3/4. A) Clusters from differential motif analysis using Gimmemotifs on formative enriched MLL3/4 independent and dependent enhancers marked by both H3K4me1 and H3K27ac (Sites from Figure 4C and 4D). Motifs derived from JASPAR 2020 database, k-means clustering used. B) Same as A but with naive enriched independent and dependent enhancers (using sites from Figure 4B and Figure S5B).

**Figure S6:**
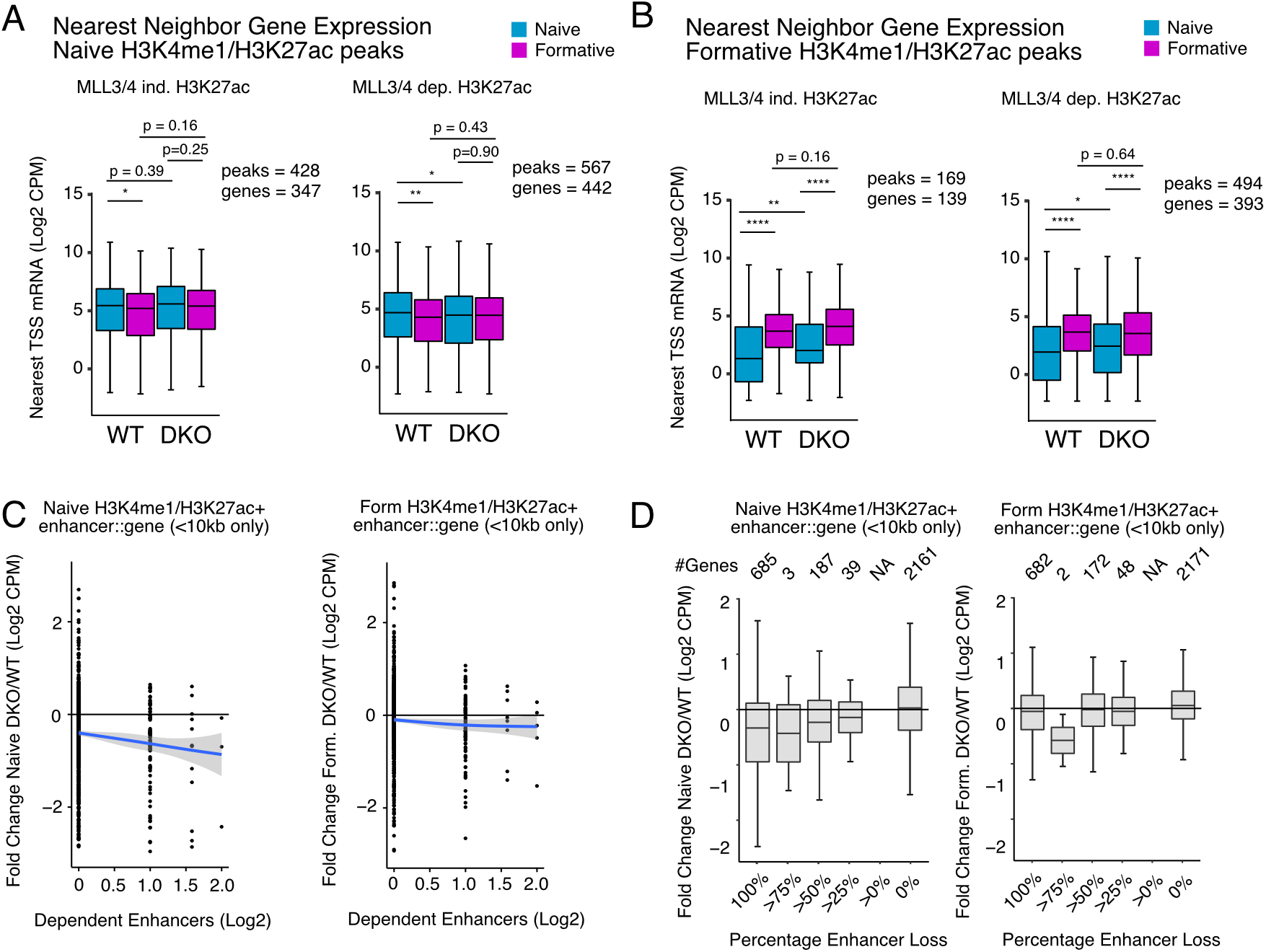
Distal H3K4me1 and H3K27ac are not functionally coupled with formative transcriptional activation. A) Nearest neighbor TSS analysis of expression levels in Log2CPM near naive H3K4me1 dependent sites that are either have MLL3/4 independent (left panel) or dependent H3K27ac (right panel)(clusters from Fig.4B). Multi-comparison paired Wilcoxon Rank-Sum Test, Benjamini-Hochberg corrected. B) Same as A but at formative H3K4me1 dependent sites (clusters from Fig.4C). C) The relative expression DKO/WT of RNA levels for all genes associated with any H3K4me1/H3K27ac+ peak in either naive or formative state compared with the number of dependent enhancers for each gene. Only dependent enhancers within 10kb of the nearest TSS are considered. Each dot represents one gene. Blue line represents generalized linear model, gray 95% confidence interval. D) Boxplots of relative expression DKO/WT of RNA levels for all genes associated with any H3K4me1/H3K27ac+ peak in either the naïve (left) or formative state (right). Each gene is binned by the percentage of their associated enhancer loss in DKOs using only enhancers within 10kb of the nearest TSS. *p<0.05 **p<0.01 ***p<0.001 ****p<0.0001

**Figure S7:**
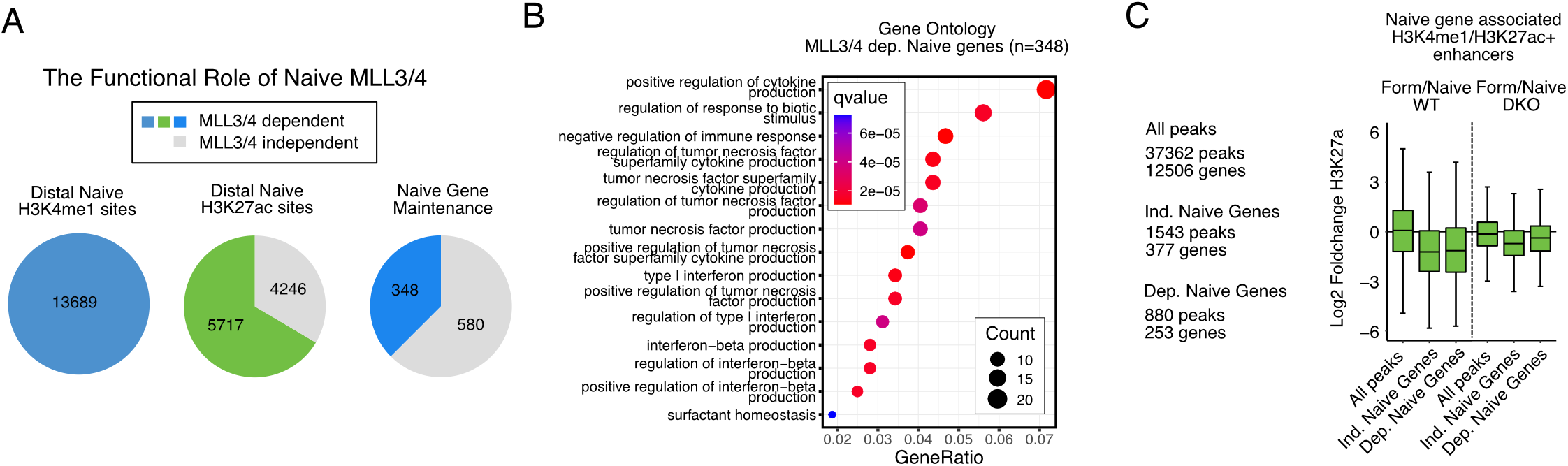
Gene-centric analysis reveals a subset of distal loci associate with MLL3/4 dependent formative genes. A) A major percentage of naive enriched peaks fail to maintain distal H3K4me1 and H3K27ac with moderate consequences on naive gene expression. B) Clusterprofile analysis of Biological Processes Gene Ontology for 348 MLL3/4 dependent naive genes. C) Fold change Log2CPM of H3K27ac density for H3K4me1/H3K27ac+ enhancers that are associated with naive genes that gained expression during transition in an either MLL3/4 independent or dependent fashion.

